# Precision RNAi in Tomato Using Synthetic Trans-Acting Small Interfering RNAs Derived From Minimal Precursors

**DOI:** 10.1101/2025.04.10.648111

**Authors:** Ariel H. Tomassi, María Juárez-Molina, Adriana E Cisneros, Ana Alarcia, Francesca Orlando, Sara Toledano-Franco, Silvia Presa, Antonio Granell, Alberto Carbonell

## Abstract

RNA interference (RNAi) is a highly conserved gene silencing mechanism regulating gene expression at transcriptional and post-transcriptional levels in plants. Synthetic trans-acting small interfering RNAs (syn-tasiRNAs) have emerged as powerful tools for highly specific and efficient gene silencing. However, their application in crops has been constrained by the need for transgene integration and the relatively long length of *TAS*-derived precursors. Here, we developed a novel syn-tasiRNA platform for *Solanum lycopersicum* (tomato) based on minimal precursors targeted by endogenous SlmiR482b microRNA. These minimal precursors, comprising only a 22-nt miRNA target site, an 11-nt spacer and the syn-tasiRNA sequence(s), effectively produced functional syn-tasiRNAs in both transgenic and transient virus-induced gene silencing (syn-tasiR-VIGS) systems. To facilitate their broader application, we engineered a series of vectors for high-throughput cloning and efficient syn-tasiRNA expression from SlmiR482b-based minimal precursors in tomato. Our results show that minimal precursors induce robust gene silencing of endogenous tomato genes and confer antiviral resistance to the economically important tomato spotted wilt virus. Furthermore, we show that syn-tasiR-VIGS can be applied in a transgene-free manner through crude extract delivery, leading to efficient silencing of endogenous genes. This study establishes minimal syn-tasiRNA precursors as a versatile and efficient tool for precision RNAi in tomato, with applications in functional genomics and crop improvement.

## INTRODUCTION

RNA interference (RNAi) is an evolutionary conserved mechanism that regulates gene expression through the sequence-specific degradation or translational repression of target messenger RNAs (mRNAs) by complementary 20-to 24-nucleotide (nt) small RNA (sRNA) molecules (Fire *et al*., 1998; Hannon, 2002). RNAi is initiated by the processing of double-stranded RNA (dsRNA) into small interfering RNAs (siRNAs) or microRNAs (miRNAs) by Dicer ribonucleases (Bernstein *et al*., 2001; Hammond *et al*., 2000). These sRNAs are subsequently incorporated into an ARGONAUTE (AGO)-containing RNA-induced silencing complex (RISC), which recognizes complementary mRNA sequences and directs their cleavage or translational suppression (Bartel, 2004; Bernstein *et al*., 2001). In plants, RNAi plays fundamental roles in endogenous gene regulation, antiviral immunity, transposon silencing and defense against pathogens (Baulcombe, 2004; Ding and Voinnet, 2007).

Classic RNAi-based approaches in plants involve the expression of dsRNA or hairpin RNA (hpRNA) transgenes to generate siRNAs homologous to the target gene(s) or the use of virus-induced gene silencing (VIGS) vectors. While effective, these methods generate heterogeneous siRNA populations, increasing the likelihood of off-target effects (Jackson *et al*., 2003). More recently, second-generation RNAi strategies based on 21-nt artificial sRNAs (art-sRNAs) were developed to improve specificity. Art-sRNAs are computationally designed to selectively target and cleave RNA transcripts with high specificity while minimizing off-target interactions (Carbonell, 2017). Art-sRNAs are categorized into two major classes: artificial microRNAs (amiRNAs) and synthetic trans-acting small interfering RNAs (syn-tasiRNAs), which operate through similar mechanisms but differ in their biogenesis. AmiRNAs are generated from *MIR* transgenes in which endogenous miRNA and miRNA* sequences are replaced with engineered amiRNA and amiRNA* sequences. Conversely, syn-tasiRNAs are derived from *TAS* transgenes, allowing the simultaneous production of multiple syn-tasiRNAs from a single precursor, facilitating multi-site targeting within one or more transcripts. Briefly, the endogenous tasiRNA sequences of *TAS* transgenes are replaced with one or more 21-nt syn-tasiRNA sequences arranged in tandem, each designed to target specific genes of interest (Zhang, 2014). *TAS* transgenes are transcribed in the nucleus by DNA-DEPENDENT RNA POLYMERASE II, producing a primary *TAS* transcript (*TAS* precursor) that contains a 5’ cap, a polyadenylated tail and a miRNA-specific target site (TS), typically 22-nt in length (Chen *et al*., 2010; Cuperus *et al*., 2010). Once exported to the cytoplasm, the *TAS* precursor is cleaved by a 22-nt miRNA/AGO complex, generating a fragment stabilized by SUPPRESSOR OF GENE SILENCING 3 (SGS3), preventing its degradation and allowing RNA-DEPENDENT RNA POLYMERASE 6 (RDR6) to synthesize a complementary dsRNA (Allen *et al*., 2005; Yoshikawa *et al*., 2005). This dsRNA is then processed into phased 21-nt syn-tasiRNA duplexes by DCL4, following a pathway similar to that of endogenous tasiRNAs. Finally, HUA ENHANCER1 (HEN1) methylates the 3’ ends of both strands, stabilizing them before the guide strand is incorporated into AGO1 (Li *et al*., 2005).

Despite their potential in functional genomics and crop improvement, syn-tasiRNA applications have been constrained by the need for *TAS*-based transgene integration, which is labor-intensive and raises regulatory concerns associated with genetically modified plants (Su *et al*., 2023). Furthermore, the relatively long length of *TAS*-derived precursors can increase RNA synthesis costs and reduces stability in viral vectors, as observed for full-length *MIR390a* and *TAS1c* amiRNA and syn-tasiRNA precursors, respectively, in *Arabidopsis thaliana* (Arabidopsis) (Cisneros, Martín-García, *et al*., 2023; Cisneros *et al*., 2025). These limitations have been recently overcome by the development of non-*TAS* syn-tasiRNA precursors of minimal length, termed “minimal” precursors, consisting of a 22-nt endogenous miRNA TS, an 11-nt spacer and the 21-nt syn-tasiRNA sequence(s). These minimal precursors were shown to generate functional syn-tasiRNAs and induce effective gene silencing when stably expressed in transgenic Arabidopsis or transiently expressed in *Nicotiana benthamiana* (Cisneros *et al*., 2025). Remarkably, they also produced authentic syn-tasiRNAs and induced widespread gene silencing in *N. benthamiana* when expressed from an RNA virus, a strategy named syn-tasiR-VIGS (Cisneros *et al*., 2025). However, whether syn-tasiRNAs can be efficiently processed and applied for stable and transgene-free gene silencing in crop species remains an open question.

In this study, we developed a syn-tasiRNA system for *Solanum lycopersicum* (tomato) using minimal precursors incorporating a SlmiR482b TS followed by an 11-nt spacer and the 21-nt syn-tasiRNA sequence(s). We show that authentic and highly effective syn-tasiRNAs can be produced from these minimal, non-*TAS* precursors in both transgenic plants and syn-tasiR-VIGS systems. Furthermore, minimal precursors successfully silenced endogenous genes and induced antiviral resistance against tomato spotted wilt virus (TSWV). Finally, we show that transgene-free syn-tasiR-VIGS, when delivered as crude extracts, resulted in efficient silencing of tomato genes. Collectively, these results establish syn-tasiRNAs as a powerful and versatile tool for gene silencing in tomato.

## RESULTS

### Syn-tasiRNA production in *S. lycopersicum* from minimal precursors using endogenous 22-nt miRNA triggers

To develop a functional syn-tasiRNA platform in *S. lycopersicum*, we first identified endogenous 22-nt miRNAs capable of triggering secondary siRNA production. We focused on SlmiR482b and SlmiR6020, both of which target nucleotide-binding leucine-rich repeat (NLR) immune receptor genes and initiate phased secondary siRNA biogenesis from target transcripts (P. Deng *et al*., 2018; Shivaprasad *et al*., 2012). Next, to assess syn-tasiRNA production in tomato, the *35S:SlmiR482bTS-NbSu* and *35S:SlmiR6020TS-NbSu* constructs were generated for expressing syn-tasiR-NbSu –a syn-tasiRNA targeting the *Nicotiana benthamiana SULPHUR* (*NbSu*) gene that accumulates to high levels in plant tissues (Cisneros *et al*., 2022) from minimal precursors including SlmiR482b or SlmiR6020 TS, respectively (Figure 1A). Agroinfiltration of these constructs was performed in separate areas of two leaves from three different tomato plants. Negative control constructs *35S:SlmiR482b-GUS*_*Nb*_ and *35S:SlmiR6020-GUS*_*Nb*_, were designed to produce syn-tasiR-GUS_Nb_, a syn-tasiRNA targeting *Escherichia coli uid*A β-glucuronidase gene (or *GUS*), from minimal precursors targeted by SlmiR482b and SlmiR6020, respectively (Figure 1A). As a positive control, the *35S:AtTAS1c-NbSu/AtMIR173A* construct was used, as it is known to efficiently produce *AtTAS1c*-based syn-tasiRNAs in tomato due to the co-expression of *Arabidopsis thaliana* AtmiR173a 22-nt miRNA (López-Dolz *et al*., 2020).

**Figure 1.**
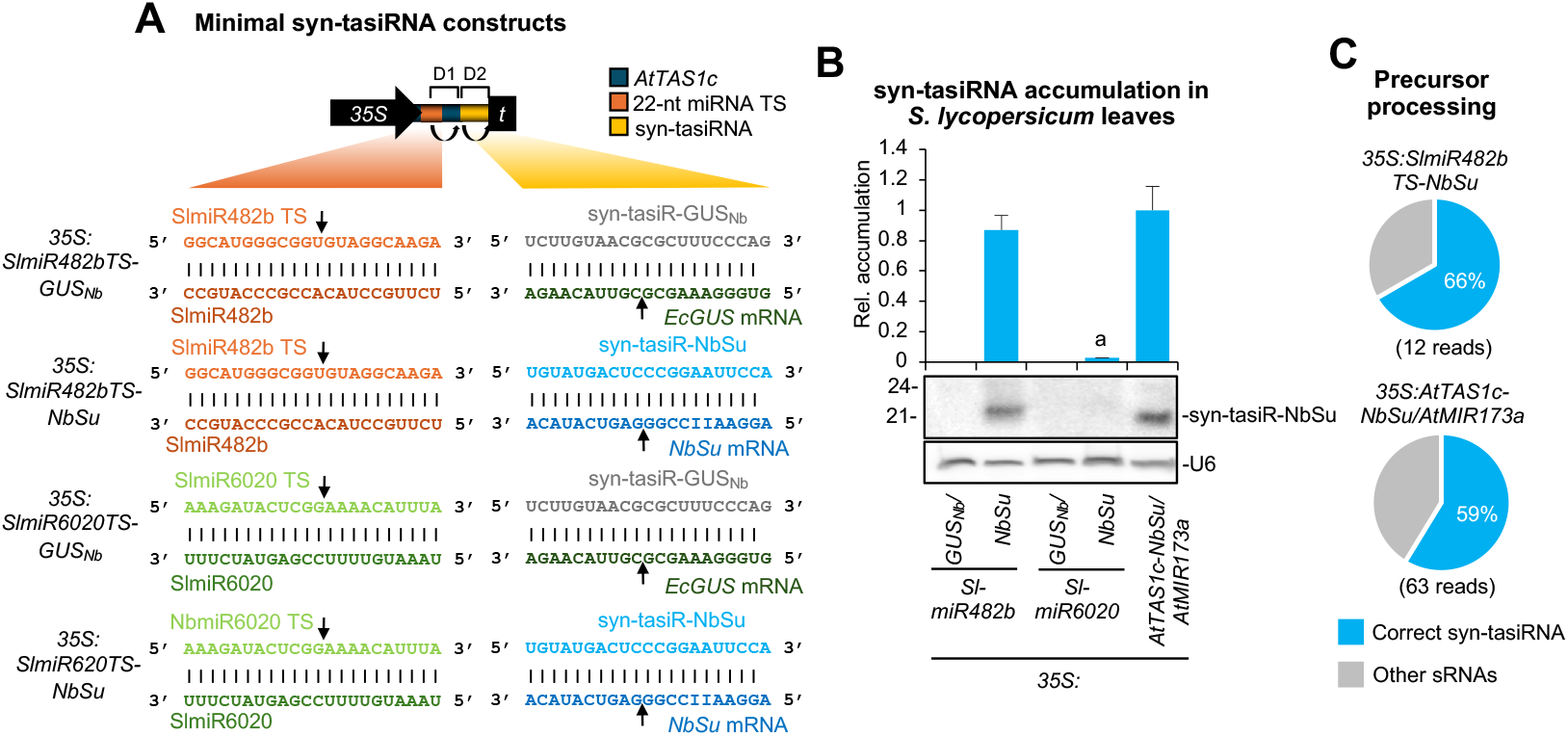
Accumulation in *Solanum lycopersicum* leaves of syn-tasiRNAs expressed from minimal precursors including endogenous 22-nt miRNA target sites (TS). (**A**) Organization of minimal precursor constructs. Nucleotides (nt) corresponding to SlmiR482b and SlmiR6020 TSs are shown in light orange and green, respectively. Nucleotides corresponding to SlmiR482b and SlmiR6020 are shown in dark orange and green, respectively. Nucleotides corresponding to syn-tasiR-GUS_Nb_ and syn-tasiR-NbSu are shown in light grey and blue, while those of their target mRNAs are shown in dark grey and blue, respectively. Arrows indicate the predicted cleavage sites for endogenous 22-nt miRNAs and syn-tasiRNAs. (**B**) Northern blot detection of syn-tasiR-NbSu in RNA preparations from agroinfiltrated leaves collected 2 days post-agroinfiltration. The graph at top shows the mean + standard deviation (n = 3) syn-tasiRNA relative accumulation (*35S:AtTAS1c-NbSu/MIR173* = 1). Bar with the letter “a” is significantly different from that of *35S:AtTAS1c-NbSu/MIR173* control samples. Each biological replicate is a pool of six agroinfiltated leaves. One blot from three biological replicates is shown. U6 blot is shown as loading control. (**C**) Syn-tasiRNA processing from mimimal precursors and controls. Pie charts show the percentages of reads corresponding to expected, accurately processed 21-nt mature syn-tasiR-NbSu (in blue) or to other 19-24 nts sRNAs (in grey).

RNA blot analysis of agroinfiltrated leaf samples collected at 2 days post-agroinfiltration (dpa) revealed high syn-tasiR-NbSu accumulation in samples expressing the *SlmiR482bTS*-based precursor, comparable to levels observed in positive control samples co-expressing *AtTAS1c-NbSu* and *AtMIR173a* (Figure 1B). In contrast, syn-tasiR-NbSu accumulated to barely detectable levels in samples expressing *SlmiR6020TS-*based minimal precursors, and, as expected, no syn-tasiR-NbSu was detected in negative control samples (Figure 1B). High-throughput sequencing of sRNAs from *35S:SlmiR482bTS-NbSu* and *35S:AtTAS1c-NbSu/AtMIR173a* samples confirmed that both precursors were processed with similar accuracy. The majority of reads within ±4 nt of the 3’D2[+] position (66% and 59%, respectively) corresponded to authentic syn-tasiR-NbSu (Figure 1C). These results demonstrate that SlmiR482b can efficiently trigger syn-tasiRNA production in tomato from minimal precursors, providing a basis for the development of an optimized syn-tasiRNA system for this species.

### High-throughput B/c vectors for syn-tasiRNA expression in tomato

To facilitate high-throughput cloning and expression of syn-tasiRNAs in tomato, we developed two new “B/c” vectors incorporating SlmiR482b: i) *pENTR-SlmiR482bTS-B/c*, a Gateway-compatible entry vector allowing the direct insertion of syn-tasiRNA sequences and subsequent recombination into preferred expression vectors with customizable promoters, terminators and regulatory elements, and ii) *pMDC32B-SlmiR482bTS-B/c*, a binary vector for direct transformation, eliminating the need for intermediate subcloning steps (Figure 2A). Both vectors contain the SlmiR482b TS followed by an 11-nt spacer sequence derived from *AtTAS1c* and a 1461 bp DNA cassette encoding the control of cell death (*ccd*B) gene (Bernard and Couturier, 1992), flanked by two inverted *Bsa*I restriction sites downstream of the 3’D1[+] position (Figure 2A). Syn-tasiRNA constructs were generated using a simplified and cost-effective cloning method, following established protocols for B/c-based vectors (Carbonell *et al*., 2014; Cisneros *et al*., 2025). Briefly, syn-tasiRNA inserts were generated by annealing two 25-nt overlapping and partially complementary oligonucleotides containing the syn-tasiRNA sequence, with 5’-TTTA and 5’-CCGA overhangs. These were then directionally ligated into *Bsa*I-digested *SlmiR482bTS-B/c* vectors (Supplementary Figure S1 and Protocol S1). The configuration of *SlmiR482bTS*-based syn-tasiRNA constructs expressing a single syn-tasiRNA is illustrated in Figure 2B. These vectors were subsequently used throughout the study to evaluate gene silencing efficacy of syn-tasiRNAs expressed from *SlmiR482bTS*-based minimal precursors.

**Figure 2.**
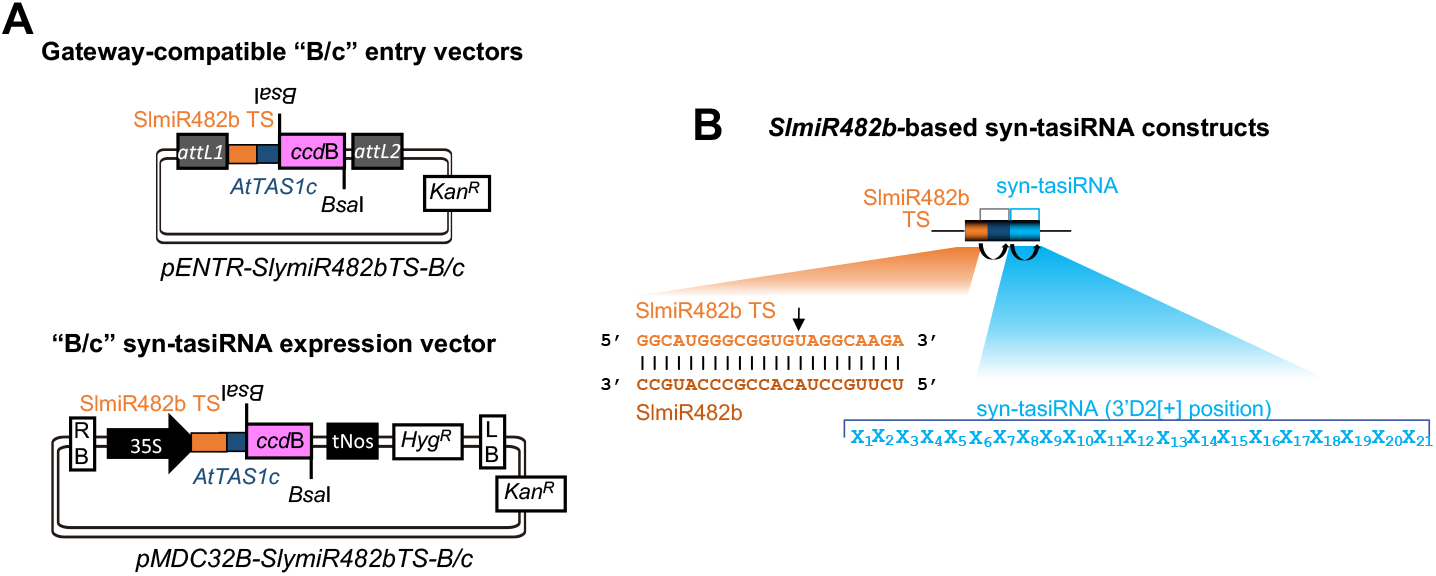
*B/c*-based vectors for direct cloning of syn-tasiRNAs downstream SlmiR482b target site (TS). (**A**) TS is in orange, the spacer sequence derived from *AtTAS1c* is in blue. Top, diagram of the Gateway-compatible entry vectors. Bottom, diagram of binary vector for in plant expression of syn-tasiRNAs. RB: right border; 35S: Cauliflower mosaic virus promoter; *Bsa*I: *Bsa*I recognition site, *ccd*B: gene encoding the gyrase toxin, in pink; LB: left border; attL1 and attL2: GATEWAY recombination sites. *Kan*^*R*^: kanamycin resistance gene; *Hyg*^*R*^: hygromycin resistance gene. (**B**) Organization of SlmiR482b-based syn-tasiRNA constructs. Base pairing between SlmiR482b (dark orange) and its target site (light orange) nucleotides is shown. Curved black arrows indicate DCL4 processing sites. Black linear arrows indicate sRNA-guided cleavage sites. TS refers to target site. In the example diagram, one single 21-nt guide syn-tasiRNA sequence was introduced at the 3’D2[+] position in *SlmiR482bTS*-based precursors.

### Generation of delayed-flowering transgenic tomato plants expressing a syn-tasiRNA against *SlSFT*

To determine whether syn-tasiRNAs expressed from *SlmiR482bTS*-based precursors could efficiently downregulate endogenous genes in tomato, we targeted the *SINGLE FLOWER TRUSS* (*SlSFT*) gene, a key regulator of flowering time (Krieger *et al*., 2010). The 21-nt syn-tasiR-SlSFT sequence, previously shown to silence *SlSFT* when expressed as an artificial miRNA (Jiang *et al*., 2013; Shalit *et al*., 2009), was cloned into the *pMDC32B-SlmiR482bTS-B/c* vector. The resulting construct, *35S:SlmiR482bTS-SlSFT* (Figure 3A), was introduced into tomato cotyledon explants via *Agrobacterium*-mediated transformation. Importantly, successful expression of syn-tasiR-SlSFT in transgenic plants was expected to effectively downregulate *SlSFT*, resulting in a delayed flowering phenotype. This trait holds significant agronomic potential, as it could extend the fruit-setting period, synchronize production with market demand and provide an extended window for crossing in breeding programs.

**Figure 3.**
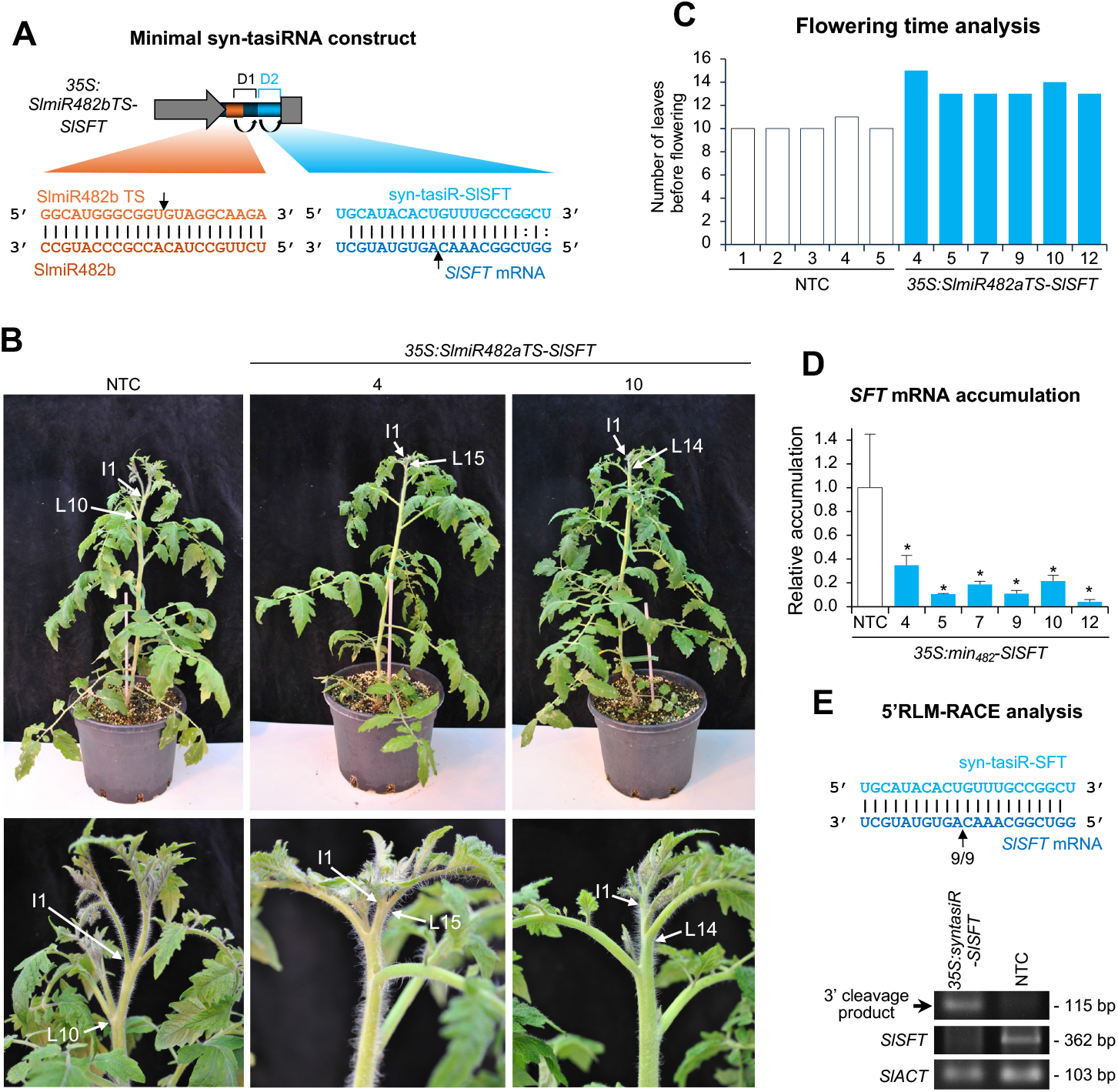
Functional analysis of *Solanum lycopersicum* T1 transgenic lines expressing syn-tasiR-SlSFT, a syn-tasiRNA targeting *SINGLE FLOWER TRUSS* (*SlSFT*). (**A**) Schematic representation of the anti-*SlSFT* syn-tasiRNA construct, *35S:SlmiR482bTS-SlSFT*, engineered to express syn-tasiR-SlSFT (light blue) from a minimal precursor containing the SlmiR482b target site (TS) (orange) and a 11-nt spacer derived from *AtTAS1c* (dark blue). Other details are as described in Figure 1A. (**B**) Photographs taken at 5 weeks post-transplanting (wpt) of representative transgenic tomato lines expressing anti-*SlSFT* syn-tasiRNA compared to a non-transgenic control (NTC) plant. Top panel: whole plants. Bottom panel: detail of the apical region, with arrows marking the first emerging inflorescence (I1) and the last leaf (L), numbered, before it. (**C**) Phenotypic analysis of flowering time in NTC and syn-tasiRNA transgenic lines, showing the number of leaves present at the time of the first emerging inflorescence in each plant. (**D**) Accumulation of *SlSFT* mRNA in tomato plants. Data are presented as the mean +SE relative expression levels of *SlSFT* mRNA at 12 wpt after normalization to *ACTIN* (SlACT), as determined by RT–qPCR (NTC=1 in all comparisons). The asterisk indicates a significant difference from the NTC samples (P<0.05; pairwise Student’s *t*-test comparison). The NTC sample corresponds to a pooled sample from five NTCs. (**E**) 5′-RLM-RACE analysis of syn-tasiR-SlSFT-guided cleavage of *SlSFT*. Upper panel: predicted base pairing between syn-tasiR-SlSFT and *SlSFT* mRNA, with the expected cleavage site indicated by an arrow. The proportion of cloned 5′-RLM-RACE products at the expected cleavage site is shown for syn-tasiR-SlSFT-expressing lines. Lower panel: ethidium bromide-stained gel showing 5′-RLM-RACE products corresponding to the 3′ cleavage product from syn-tasiR-SlSFT-guided cleavage (top), along with RT– PCR products for the target *SlSFT* (middle) and the control *SlACT* genes (bottom). The position and expected sizes of syn-tasiRNA-based 5′-RLM-RACE products and control RT-PCR products are indicated.

Phenotypic analysis of six independent T1 transgenic lines at 5 weeks post-transplanting (wpt) revealed a significant delay in flowering compared to non-transgenic control (NTC) plants (Figure 3B). While NTC plants produced an average of 10 leaves before the emergence of the first inflorescence (I1), syn-tasiR-SlSFT-expressing transgenic lines exhibited an increased number of leaves (13–15) before I1 formation (Figure 3C), confirming a delayed transition from the vegetative to the reproductive phase. To assess the molecular effects of syn-tasiR-SlSFT, RT–qPCR analysis was performed at 12 wpt to measure *SlSFT* transcript levels. Our results show a significant reduction in *SlSFT* mRNA accumulation in syn-tasiR-SlSFT-expressing plants compared to NTCs (Figure 3D), indicating effective suppression of *SlSFT* expression. To confirm that the reduced *SlSFT* transcript levels were due to syn-tasiR-SlSFT-mediated cleavage, 5′-RLM-RACE analysis was conducted. Gel electrophoresis confirmed the amplification of the expected 3′ cleavage fragments in transgenic plants, whereas no cleavage products were detected in NTC samples (Figure 3E, lower panel). Sequence analysis of 5’-RLM-RACE products revealed precise cleavage at the expected target site within the *SlSFT* mRNA, with a high proportion of cleavage events occurring at the predicted position (Figure 3E, upper panel). RT-PCR analysis further confirmed the downregulation of *SlSFT* in transgenic plants, while *SlACT* expression was used as a reference to validate RNA integrity and cDNA synthesis across all samples. Collectively, these results demonstrate that syn-tasiR-SlSFT, produced from *SlmiR482bTS*-based minimal precursors, effectively downregulates *SlSFT* in transgenic tomato plants, leading to a prolonged vegetative phase and a delayed flowering.

Next, to assess the accumulation and processing efficiency of syn-tasiR-SlSFT, northern blot analysis of RNA preparations from apical leaves at 12 wpt confirmed the presence of a 21-nt RNA species corresponding to syn-tasiR-SlSFT in transgenic lines, whereas no detectable signal was observed in NTC plants (Figure 4A). High-throughput sRNA sequencing further confirmed that syn-tasiR-SlSFT was accurately processed. Analysis of 19-to 24-nt reads mapping to *SlmiR482bTS-SlSFT* precursors revealed a major peak corresponding of 21-nt reads starting at the expected syn-tasiRNA position (position 34) and corresponding to syn-tasiR-SlSFT (Figure 4B). Additionally, 87% of reads within ±4 nt of the 3’D2[+] position corresponded to authentic syn-tasiR-SlSFT, with only a minor fraction mapping to other regions (Figure 4C), indicating that *SlmiR482bTS*-based precursors are efficiently recognized and cleaved in tomato. To further evaluate the phased processing of the syn-tasiRNA precursor, we analyzed the register distribution of 21-nt reads along the precursor transcript. A radar plot showed that 98% of 21-nt reads aligned to the expected register, confirming that tasiRNA processing occurs with high phasing precision in tomato (Figure 4D). Finally, the absence of 21-nt siRNAs derived from *SlSFT* (Figure S2, Data S1) indicates that syn-tasiR-SlSFT is only causing the intended cleavage of *SlSFT*, without triggering an amplification loop of secondary, 21-nt siRNAs that could target other regions of the mRNA or similar sequences. Together, these results confirm that syn-tasiR-SlSFT is stably expressed, accurately processed from *SlmiR482bTS*-based minimal precursors, and induces specific and robust gene silencing in tomato.

**Figure 4.**
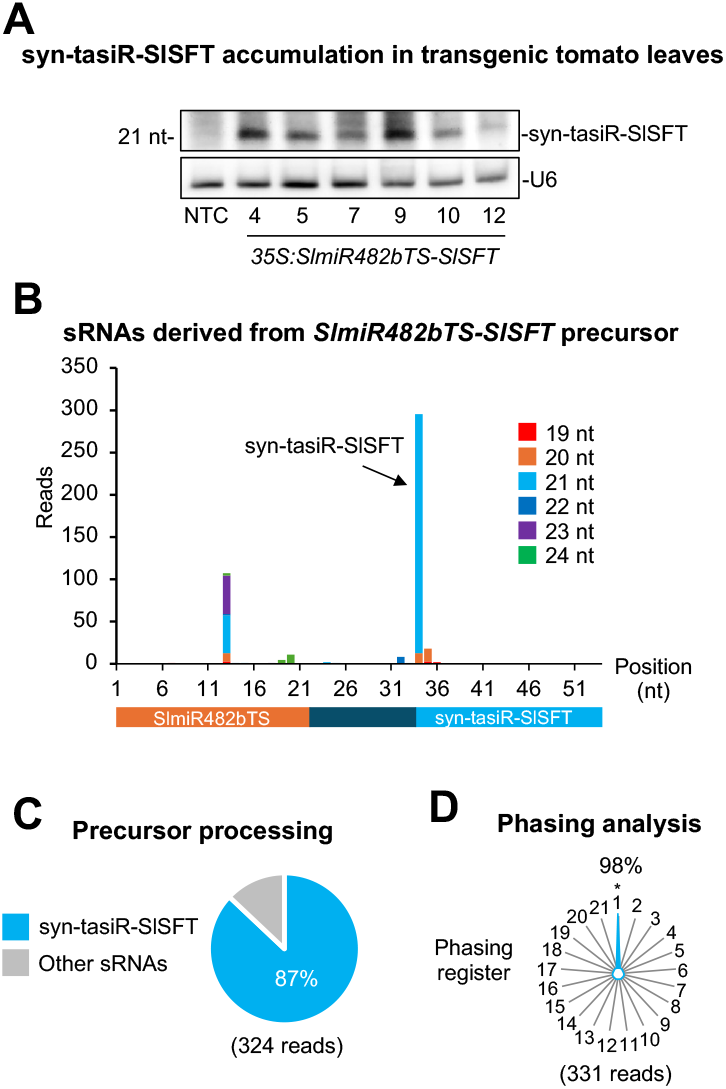
Accumulation and processing of syn-tasiR-SlSFT expressed from *SlmiR482bTS*-based precursors in *Solanum lycopersicum* T1 transgenic lines. (**A**) Northern blot detection of syn-tasiR-SlSFT in RNA preparations from apical leaves collected 12 weeks post-transplanting from six independent transgenic lines and one non-transgenic control (NTC). Each sample represents a pool of two apical leaves. Other details are as described in Figure 1B. (**B**) sRNA profile of 19-24 nt [+] reads mapping to each of the 54 nucleotide positions within the *SlmiR482bTS-SlSFT* precursor from plants expressing *35S:SlmiR482bTS-SlSFT*. Orange, dark blue and light blue boxes represent nucleotides corresponding to *NbmiR482aTS*, the *AtTAS1c*-derived spacer and syn-tasiR-SlSFT, respectively. (**C**) Pie chart showing the percentage of reads corresponding to accurately processed 21-nt authentic syn-tasiR-SlSFT (blue) versus other 19-24-nt sRNAs (gray). (**D**) Radar plot displaying the distribution of 21-nt reads across the 21 registers of the precursor transcripts, with position 1 designated immediately after the SlmiR482b-guided cleavage site.

### Syn-tasiR-VIGS antiviral vaccination of tomato plants against TSWV

Next, to evaluate the potential of using a viral vector for syn-tasiRNA-mediated gene silencing in tomato, we investigated the efficacy of potato virus X (PVX)-based syn-tasiR-VIGS to confer resistance against tomato spotted wilt virus (TSWV). Several PVX-based constructs were generated for expressing syn-tasiRNAs from minimal precursors (Figure 5A). The *35S:PVX-SlmiR482bTS-TSWV(x4)* construct included the *SlmiR482bTS* precursor engineered to produce four highly active syn-tasiRNAs (syn-tasiR-TSWV-1 to syn-tasiR-TSWV-4) targeting conserved regions of TSWV genome (Carbonell, Lisón, *et al*., 2019; Carbonell, Lopez, *et al*., 2019) (Figure 5B). Negative control constructs included the *35S:PVX-SlmiR482bTS-GUS*_*Sl*_*(x4)* construct, designed to express two syn-tasiRNAs (syn-tasiR-GUS_Sl_-1 and syn-tasiR-GUS_Sl_-2) against *GUS* (with no off-targets in tomato) (Carbonell, Lisón, *et al*., 2019) from *SlmiR482bTS*-based precursors, and *35S:PVX-AtmiR173aTS-TSWV(x4)*, including the AtmiR173a TS, which was expected to fail in triggering syn-tasiRNA biogenesis due to the absence of AtmiR173a in *S. lycopersicum* (Figure 5B), as reported before in *N. benthamiana* (Cisneros *et al*., 2025).

**Figure 5.**
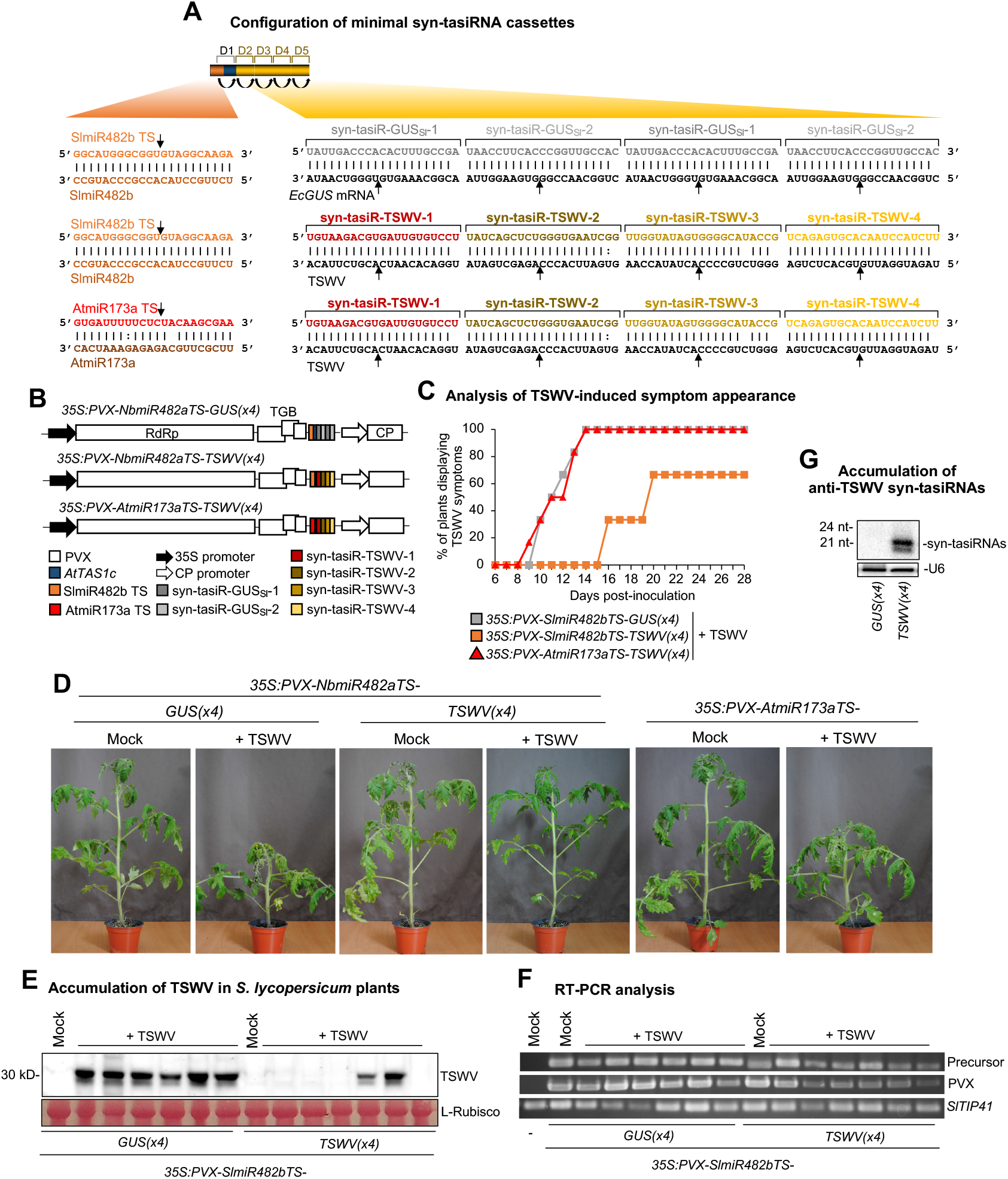
Functional analysis of potato virus X (PVX) constructs expressing syn-tasiRNAs against tomato spotted wilt virus (TSWV) in *S. lycopersicum*. (**A**) Schematic representation of PVX-based constructs. Nucleotides of anti-TSWV art-sRNA sequences syn-tasiR-TSWV-1, syn-tasiR-TSWV-2, syn-tasiR-TSWV-3 and syn-tasiR-TSWV-4 are in red, dark brown, light brown and yellow, respectively. Nucleotides of control anti-GUS art-sRNA sequences syn-tasiR-GUS_Sl_-1 and syn-tasiR-GUS_Sl_-2 are in dark and light grey, respectively. Nucleotides of AtmiR173a target site (TS) are in red and brown, respectively. Other details are as in Figure 1A. (**B**) Diagram of PVX-based constructs expressing anti-TSWV or anti-*GUS* syn-tasiRNAs. Color coding for the syn-tasiRNA sequences is consistent with panel (A). Other details are as in Figure 1A. (**C**) Two-dimensional line graph showing, for each of the six-plant sets listed, the percentage of symptomatic plants per day during 28 days. (**D**) Photographs taken at 14 days post-inoculation (dpi) of plants agroinoculated with the different constructs and inoculated (+TSWV) or not (mock) with TSWV. (**E**) Western blot detection of TSWV in protein extracts from apical leaves collected at 14 dpi. A Ponceau-stained membrane is shown as a loading control, highlighting the large subunit of Rubisco (ribulose1,5-biphosphate carboxylase/oxygenase). (**F**) RT-PCR detection at 14 dpi of *SlmiR482bTS*-based precursors and PVX coat protein fragment (PVX-CP) in apical leaves agroinoculated plants. RT-PCR amplification of *SlTYP41* is included as a control. (**G**) Northern blot detection of anti-TSWV art-sRNAs in RNA preparations from apical leaves collected at 14 dpi. A cocktail of probes to simultaneously detect syn-tasiR-TSWV-1, syn-tasiR-TSWV-2, syn-tasiR-TSWV-3 and syn-tasiR-TSWV-4 was used. Other details are as in Figure 1C.

To assess the antiviral activity of PVX-based constructs, each construct was agroinoculated into two leaves of six independent plants. Twelve days later, these same plants were further inoculated with TSWV, and symptom progression was monitored over 28 days. At 14 days post inoculation (dpi), all plants expressing *35S:PVX-SlmiR482bTS-TSWV(x4)* remained asymptomatic, whereas control plants agroinoculated with *35S:PVX-SlmiR482bTS-GUS*_*Sl*_*(x4)* or *35S:PVX-AtmiR173aTS-TSWV(x4)* exhibited typical TSWV symptoms (Figure 5C and 5D), which developed and progressed similarly over time. Since plants from the two control blocks exhibited similar behavior, subsequent molecular analyses were conducted comparing plants agroinoculated with *35S:PVX-SlmiR482bTS-TSWV(x4)* to those agroinoculated with *35S:PVX-SlmiR482bTS-GUS*_*Sl*_*(x4)*. At 14 dpi, western blot analysis revealed that four out of six plants expressing *35S:SlmiR482bTS-TSWV(x4)* did not accumulate detectable TSWV, while plants expressing *35S:SlmiR482bTS-GUS*_*Sl*_*(x4)* accumulated high TSWV levels (Figure 5E). By 28 dpi, two plants expressing anti-TSWV syn-tasiRNAs from *SlmiR482bTS*-based precursors remained completely symptom-free (Figure 5C). Importantly, RT-PCR analysis confirmed the presence of a 341-bp fragment corresponding to the *SlmiR482bTS*-based precursors and a 230-bp fragment from the PVX coat protein (CP) in all PVX-treated samples, but not in mock-inoculated and non-agroinfiltrated plants, while a 235 bp fragment from *SlTIP41* was amplified in all samples (Figure 5F). Finally, northern blot analysis at 14 dpi confirmed the presence of high levels of 21-nt anti-TSWV syn-tasiRNAs in transgenic plants expressing *35S:SlmiR482bTS-TSWV(x4)*, whereas no corresponding signals were detected in *35S:PVX-SlmiR482bTS-GUS*_*Sl*_*(x4)* control samples (Figure 5G). These results indicate that syn-tasiRNA precursors were accurately processed and accumulated to levels sufficient for antiviral activity. Collectively, these results indicate that tomato vaccination with syn-tasiR-VIGS interferes with TSWV infection and can confer durable antiviral resistance.

### Transgene-free syn-tasiR-VIGS in tomato

Next, we investigated the possibility of applying PVX-based syn-tasiR-VIGS in tomato in a transgene-free manner. Using P-SAMS (Fahlgren *et al*., 2016), we designed two highly specific syn-tasiRNAs (with no off-targets in *S. lycopersicum*) targeting *SULPHUR* (*SlSu*), a key enzyme in chlorophyll biosynthesis, or *1-DEOXY-D-XYLULOSE-5-PHOSPHATE SYNTHASE* (*SlDXS*), which plays a role in plastidic isoprenoid biosynthesis (Figure 6A). Silencing either gene was expected to induce bleaching phenotypes in tomato leaves. *SlmiR482bTS*-based precursors including these syn-tasiRNAs were inserted into a PVX infectious clone to generate the *35S:PVX-SlmiR482bTS-SlSu-(x2)* and *35S:PVX-SlmiR482bTS-SlDXS(x2)* constructs (Figure 6B). The negative control *35S:PVX-SlmiR482bTS-GUS*_*Sl*_*(x2)* construct including syn-tasiR-GUS_Sl_-1 and syn-tasiR-GUS_Sl_-2 sequences in tandem was also generated (Figure 6A and 6B). To prepare crude extracts for transgene-free syn-tasiR-VIGS, *Nicotiana benthamiana* plants were agroinfiltrated with PVX-based syn-tasiRNA constructs, and apical leaves were harvested 7 dpa for crude extract preparation (Figure 6C). These extracts were subsequently sprayed onto young tomato plants to induce silencing of *SlSu* or *SlDXS* (Figure 6C).

**Figure 6.**
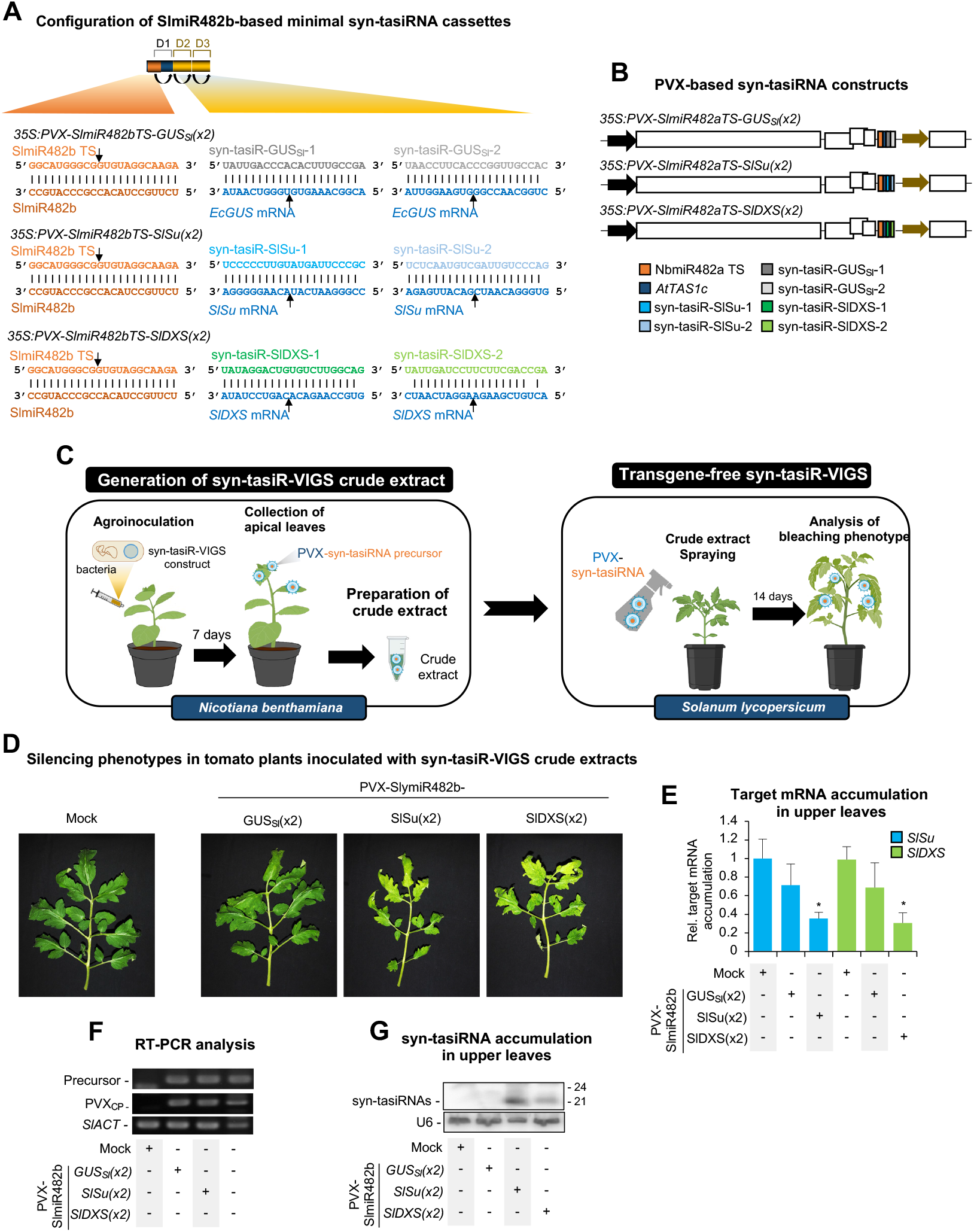
Transgene-free gene silencing through PVX-based syn-tasiR-VIGS in *Solanum lycopersicum*. (**A**) Schematic representation of PVX-based constructs. Nucleotides of anti-*SlSu* syn-tasiRNAs (syn-tasiR-SlSu-1 and syn-tasiR-SlSu-2) are shown in dark and light blue, respectively. Nucleotides of anti-*SlDXS* syn-tasiRNAs (syn-tasiR-SlDXS-1 and syn-tasiR-SlDXS-2) are shown in dark and light green, respectively. Other details are as in Figure 1A and Figure 5A. (**B**) Diagram of PVX-based constructs expressing anti-*SlSu*, anti-*SlDXS* or anti-*GUS* syn-tasiRNAs. Color coding for the syn-tasiRNA sequences is consistent with panel (A). Other details are as in Figures 1A and 5A. (**C**) Experimental procedure for transgene-free syn-tasiR-VIGS in *S. lycopersicum*. Left: crude extracts are prepared from *Nicotiana benthamiana* plants previously agroinfiltrated with the corresponding syn-tasiR-VIGS construct. Right: young tomato plants are spray-inoculated with syn-tasiR-VIGS crude extracts to induce bleaching associated with *SlSu* or *SlDXS* silencing. (**D**) Representative photographs of tomato leaves at 14 days post-spray (dps), from plants sprayed with different crude extracts obtained from agroinoculated *N. benthamiana* plants. (**E**) Accumulation of *SlSu and SlDXS* mRNA in tomato plants treated with syn-tasiR-VIGS crude extracts. Data are presented as the mean +SE relative expression levels of *SlSu or SlDXS* mRNA at 14 dps after normalization to *ACTIN* (*SlACT*), as determined by RT–qPCR (Mock=1 in all comparisons). The asterisk indicates a significant difference from the mock samples (P<0.05; pairwise Student’s *t*-test comparison). (**F**) RT-PCR detection at 14 dps of *SlmiR482bTS*-based precursors and PVX coat protein fragment (PVX-CP) in apical leaves of sprayed plants. RT-PCR amplification of *SlACT* is included as a control. (**G**) Northern blot detection of anti-*SlSu* and anti-*SlDXS* syn-tasiRNAs in RNA preparations from apical leaves collected at 14 dps. A cocktail of probes to simultaneously detect syn-tasiR-SlSu-1, syn-tasiR-SlSu-2, syn-tasiR-SlDXS-1 and syn-tasiR-SlDXS-2 was used. Other details are as described in Figure 1C.

At 14 days post-spraying (dps), tomato plants treated with crude extracts containing anti-*SlSu* or anti-*SlDXS* syn-tasiRNAs exhibited a visible bleaching phenotype in young leaves, consistent with effective gene silencing (Figure 6D). Control plants sprayed with crude extracts containing control anti-*GUS* syn-tasiRNAs showed no bleaching, further confirming the specificity of syn-tasiRNA-mediated silencing. RT-qPCR analyses on RNA preparations from upper leaves at 14 dps revealed significantly reduced *SlSu* and *SlDXS* transcript levels in plants treated with PVX-SlmiR482bTS-SlSu(x2) or PVX-SlmiR482bTS-SlDXS(x2), respectively (Figure 6E), compared to controls. At this same time point, RT-PCR analysis confirmed the presence of a 299-bp fragment corresponding to the *SlmiR482bTS*-based precursors and a 230-bp fragment from the PVX CP in all PVX-treated samples, but not in mock-inoculated plants, while a 103 bp fragment from *SlACT* was amplified in all samples (Figure 6F). Finally, northern blot analysis confirmed high accumulation of syn-tasiRNAs in these plants (Figure 6G). Taken together, these results show that syn-tasiRNAs can be efficiently produced from PVX crude extracts, accumulate to effective levels, and lead to robust and specific gene silencing in tomato in a transgene-free manner.

## DISCUSSION

In this study, we show that minimal syn-tasiRNA precursors, consisting of only a 22-nt SlmiR482b TS and an 11-nt spacer, generate high levels of accurately processed syn-tasiRNAs in *Solanum lycopersicum*, allowing efficient gene silencing in both stable and transient, transgene-free applications. Our findings establish a robust and optimized syn-tasiRNA platform in tomato, overcoming limitations associated with classic RNAi technologies and expanding the potential applications of syn-tasiRNAs in functional genomics and crop improvement.

The successful adaptation of syn-tasiRNA technology to tomato using minimal precursors targeted by endogenous 22-nt miRNAs significantly advances RNAi-based gene silencing tools for crops. Previous studies have reported the efficacy of transgenically-expressed syn-tasiRNAs derived from full-length *TAS* precursors in silencing endogenous genes in model species such as *Arabidopsis thaliana* and *N. benthamiana* (Felippes and Weigel, 2009; de la Luz Gutierrez-Nava *et al*., 2008; Montgomery, Howell, *et al*., 2008; Montgomery, Yoo, *et al*., 2008). More recently, *TAS*-based syn-tasiRNAs were used to confer antiviral resistance in tomato (Carbonell, Lisón, *et al*., 2019). However, *AtTAS1c*-based syn-tasiRNA constructs require co-expression of *AtMIR173a* in non-Arabidopsis species, adding complexity to their implementation in crops such as tomato and *N. benthamiana*. Recent research has shown that minimal syn-tasiRNA precursors targeted by 22-nt miRNAs can generate authentic syn-tasiRNAs in *N. benthamiana* (Cisneros *et al*., 2025). Here, we extended this strategy to tomato, using the endogenous 22-nt miRNAs SlmiR482b and SlmiR6020, both of which trigger secondary siRNA production from target transcripts (Y. Deng *et al*., 2018; Shivaprasad *et al*., 2012). Our results show that precursors targeted by SlmiR482b produced higher syn-tasiRNAs levels than those targeted by SlmiR6020, most likely because of the stronger expression of SlmiR482b in these samples (Figure S3). This is consistent with previous observations in *N. benthamiana*, where NbmiR482a was expressed to higher levels than NbmiR6019a/b, inducing high syn-tasiRNA production (Cisneros *et al*., 2025). Furthermore, syn-tasiRNA accumulation from full-length *AtTAS1c* precursors was comparable in leaves overexpressing AtmiR173a and in those using SlmiR482b as the trigger (Figure 1B), suggesting that SlmiR482b levels are not a limiting factor in tomato. Since miRNA expression varies across tissues, developmental stages and environmental conditions (Jones-Rhoades *et al*., 2006), selecting an optimal endogenous miRNA trigger is crucial for precise syn-tasiRNA biogenesis. Future studies could explore tissue- or stress-specific expression of endogenous miRNAs to enhance spatiotemporal control of syn-tasiRNA production in crops.

A key feature of syn-tasiRNA technology is its high specificity, as 21-nt guide sequences are computationally designed based on target specificity, minimizing off-target effects (Fahlgren *et al*., 2016; Ossowski *et al*., 2008). In contrast, larger *TAS*-based systems generate heterogenous populations of secondary siRNAs from gene fragments inserted after the 22-nt miRNA TS (Felippes and Weigel, 2009; Zhao *et al*., 2024), which can lead to unintended molecular interactions and off-target phenotypes. Here, our high-throughput sequencing data confirms that minimal precursors are accurately processed in tomato, yielding authentic and correctly phased syn-tasiRNAs that accumulate at high levels of relative to other sRNAs derived from the precursor (Figure 4B). Furthermore, 21-nt syn-tasiRNAs accumulate in plants treated with syn-tasiR-VIGS constructs or crude extracts, indicating efficient production from PVX-SlmiR482bTS-based vectors, in line with previous findings in *N. benthamiana* using PVX-NbmiR482aTS-based vectors (Cisneros *et al*., 2025). Importantly, the absence of resistance in plants expressing *35S:PVX-AtmiR173aTS-TSWV(x4)* further supports that viral protection resulted from syn-tasiRNA activity rather than from potential siRNAs generated from the minimal precursor during PVX replication.

The use of minimal syn-tasiRNA precursors offers several biotechnological advantages. First, shorter syn-tasiRNA precursors reduce construct synthesis costs, making them ideal for large-scale applications that require multiple art-sRNA constructs (Hauser *et al*., 2013; Jover-Gil *et al*., 2014; Zhang *et al*., 2018). Here, SlmiR482b-targeted precursors were integrated into a new series of *B*/c vectors optimized for cost-effective, high-throughput cloning of syn-tasiRNAs for tomato. This Gateway-compatible and direct binary vector cloning system allows rapid syn-tasiRNA deployment while simplifying construct generation, addressing one of the primary challenges in previous RNA-based technologies. In particular, our optimized *B/*c system allows for easy customization and integration of different promoter and terminator elements in the *pENTR-SlmiR482bTS-B/c* vector. Second, minimal syn-tasiRNA precursors can improve the efficiency of *in vitro* or bacterial synthesis, facilitating their large-scale production for use in topical application to plants. Third, minimal syn-tasiRNA precursors remain stable in treated tomato plants, whereas full-length *TAS*-based precursors are often lost during extended infections (Cisneros *et al*., 2025). The compact size of minimal precursors may be particularly convenient when using viral vectors with limited cargo capacity. And fourth, syn-tasiR-VIGS allows for the application of syn-tasiRNAs through crude extract spraying, providing a non-transgenic, scalable method for delivering highly specific art-sRNA molecules to plants. It is important to note that the efficacy of our syn-tasiR-VIGS approach, particularly for inducing anti-TSWV resistance, depends on selecting validated syn-tasiRNAs with proven specificity (Carbonell, Lopez, *et al*., 2019). This compensates for producing fewer but highly efficient syn-tasiRNAs, unlike larger *TAS* systems that generate uncontrolled secondary siRNAs from gene fragments, as explained before.

Despite its biotechnological potential, several aspects of syn-tasiRNA technology require further study. Currently, syn-tasiRNA minimal precursors are species-specific, the identification of an efficient endogenous 22-nt miRNA trigger for each plant species. Regarding syn-tasiR-VIGS, alternative viral vectors beyond PVX should be explored to extend syn-tasiR-VIGS to non-*Solanaceae* crops. As with all VIGS approaches, potential challenges include potential off-target effects due to viral vector infection, inconsistent efficiency depending on the plant species, tissues and environmental conditions, and vector mutations during prolonged infections that could compromise efficacy (Rössner *et al*., 2022). Finally, the integration of syn-tasiRNA technology into commercial breeding programs will require navigating regulatory frameworks for RNA-based technologies. While RNAi is generally considered a non-GMO approach, regulations vary internationally, particularly regarding the use of recombinant viral vectors for delivery. Addressing these regulatory challenges and developing clear guidelines will be critical for the commercialization of syn-tasiRNA-based products.

In conclusion, our study establishes a functional and versatile syn-tasiRNA system in tomato, providing a new platform for precision RNAi in both stable and transgene-free applications. Future research should focus on expanding syn-tasiRNA applications across multiple crops, optimizing delivery strategies, and addressing regulatory considerations to facilitate the widespread adoption of this promising RNA-based technology in agriculture.

## MATERIALS AND METHODS

### Plant species and growing conditions

*S. lycopersicum* cv Moneymaker and *N. benthamiana* plants were grown in a controlled growth chamber at 25ºC, with a 12 h light/12 h dark photoperiod. Plants were photographed using a Nikon D3000 digital camera equipped with an AF-S DX NIKKOR 18-55 mm f/3.5–5.6G VR lens.

### Generation and phenotyping of tomato cv. Moneymaker transgenic plants

*Agrobacterium tumefaciens* LBA4404 transformed with *35S:syn-tasiR-SlSFT* was co-cultured with tomato cotyledons. Explant preparation, selection and regeneration were performed as previously described (Ellul *et al*., 2003). Transformants were selected in hygromycin-containing medium, then transferred to soil for propagation, seed production and phenotypic and molecular analyses. Non-transgenic controls (NTCs) were *in vitro*-regenerated tomato plants obtained in parallel with the transgenic plants. Phenotyping was conducted using five NTCs and six independent syn-tasiRNA lines. Flowering time was assessed by counting the number of leaves at the emergence of the first inflorescence.

### Artificial small RNA design

P-SAMS script (https://github.com/carringtonlab/psams) (Fahlgren *et al*., 2016), configured to return unlimited optimal results, was used to obtain a list of optimal amiRNAs targeting *SlSu* and *SlDXS* with high specificity (Data S2). Off-target filtering was applied using the *S. lycopersicum* transcriptome iTAGv4.0 (https://solgenomics.net/ftp/tomato_genome/annotation/ITAG4.0_release/) (Hosmani *et al*., 2019) to enhance syn-tasiRNA specificity. The guide sequences for syn-tasiR-NbSu, syn-tasiR-GUS_Nb_, syn-tasiR-SlSFT, syn-tasiR-GUS_Sl_-1, syn-tasiR-GUS_Sl_-2, syn-tasiR-TSWV-1, syn-tasiR-TSWV-2, syn-tasiR-TSWV-3 and syn-tasiR-TSWV-4 were described before (Carbonell, Lopez, *et al*., 2019; Carbonell and Daròs, 2017; Cisneros *et al*., 2022; Shalit *et al*., 2009).

### DNA constructs

Oligonucleotides AC-902/AC-903 including SlmiR482b targeted precursor were annealed and ligated into *Bsa*I-digested *pENTR-B/c* and *pMDC32B-B/c* to generate *pENTR-SlmiR482bTS-BB* and *pMDC32B-SlmiR482bTS-BB*, respectively. The B/c cassette was excised from *Bsa*I-digested *pMDC32B-AtTAS1c-D2-B/c* (Addgene plasmid #137884) (López-Dolz *et al*., 2020) and ligated into *Bsa*I-digested *pENTR-SlmiR482bTS-BB* and *pMDC32B-SlmiR482bTS-BB* to generate *pENTR-SlmiR482bTS-B/c* and *pMDC32B-SlmiR482bTS-B/c*. These new B/c vectors *pENTR-SlmiR482bTS-B/c* (Addgene plasmid #234368) and *pMDC32B-SlmiR482bTS-B/c* (Addgene plasmid #234369) were deposited in Addgene.

Constructs *35S:SlmiR482b-GUS*_*Sl*_, *35S:SlmiR482b-NbSu, 35S:SlmiR6019-GUS*_*Sl*_, *35S:SlmiR6019-NbSu, 35S:SlmiR482b-SlSu(x2), 35S:SlmiR482b-SlDXS(x2), 35S:SlmiR482b-GUS*_*Sl*_*(x2)* and *35S:SlmiR482b-SlSFT* were obtained by ligating annealed oligonucleotide pairs AC-824/AC-825–AC-826/AC-827–AC828/AC-829–AC-830/AC-831–AC-892/AC-893–AC-894/AC-895–AC-915/AC-916 and AC-909/AC-910 into *pMDC32B-B/c* (Addgene plasmid #227963) (Cisneros *et al*., 2024) and *pMDC32B-SlmiR482bTS-B/c*, respectively, as described (Carbonell *et al*., 2014) (Figure S1). A detailed protocol for cloning syn-tasiRNAs in B/c vectors including SlmiR482b TS is described in Text S1.

For PVX-based constructs, syn-tasiRNA cassettes *AtmiR173aTS-TSWV(x4), SlmiR482bTS-GUS*_*Sl*_*(x2), SlmiR482bTS-SlSu(x2)* and *SlmiR482bTS-SlDXS(x2)* were synthesized as dsDNA oligonucleotides AC-1205, AC-993, AC-775 and AC-776, respectively, and assembled into *Mlu*I-digested and gel-purified *pLBPVXBa-M* (Addgene plasmid #229079) (30) in the presence of GeneArt Gibson Assembly HiFi Master Mix (Invitrogen) to generate *35S:PVX-AtmiR173aTS-TSWV(x4), 35S:PVX-SlmiR482bTS-GUS*_*Sl*_*(x2), 35S:PVX-SlmiR482bTS-SlSu(x2)* and *35S:PVX-SlmiR482bTS-SlDXS(x2)*, respectively. Text S2 provides a detailed protocol for cloning syn-tasiRNA *SlmiR482bTS*-based minimal precursors into *pLBPVXBa-M*.

*35S:AtTAS1c-NbSu/MIR173, 35S:PVX-NbmiR482aTS-GUSsl(x4) and 35S:PVX-NbmiR482aTS-TSWV(x4)* were described before (Cisneros *et al*., 2025; López-Dolz *et al*., 2020). All DNA oligonucleotides used in this study are listed in Table S1. The sequences of all syn-tasiRNA precursors are listed in Text S3. The sequences of newly developed B/c vectors are listed in Text S4.

### Transient expression of constructs and inoculation of viruses

*Agrobacterium*-mediated infiltration of DNA constructs in *N. benthamiana* and *S. lycopersicum* leaves was done as previously (Carbonell *et al*., 2015; Cuperus *et al*., 2010). Preparation of crude extracts obtained from virus infected *N. benthamiana* plants was done as previously (Cisneros, Martín-García, *et al*., 2023; Cisneros *et al*., 2025). Syn-tasiR-VIGS extracts were sprayed onto *S. lycopersicum* leaves, and TSWV was mechanically inoculated by gently rubbing the leaf surface (Carbonell, Lopez, *et al*., 2019; Cisneros *et al*., 2025).

### Total RNA preparation

Total RNA form *S. lycopersicum* leaves was isolated as previously described (Cisneros, Lisón, *et al*., 2023). Triplicate samples from pools of two leaves were analyzed.

### Real-time RT-qPCR

Real time RT-qPCR was performed using the same RNA samples as those used for sRNA-blot analysis, as described (Cisneros *et al*., 2025). Oligonucleotides used for RT-qPCR are listed in Table S1. Target mRNA expression levels were calculated relative to the reference gene *SlACT* using the delta delta cycle threshold comparative method in QuantStudio Design and Analysis software (version 1.5.1, Thermo Fisher Scientific). Three independent biological replicates were analyzed, each with two technical replicates.

### Stability of syn-tasiRNA precursors during viral infections

Total RNA from apical leaves of the three biological replicates was pooled prior cDNA synthesis. PCR detection of syn-tasiRNA precursors, PVX and *SlACT* was performed using oligonucleotide pairs AC-654/AC-655, AC-657/AC-658 and AC-280/AC-281 (Table S1), respectively, with Phusion DNA polymerase (ThermoFisher Scientific). PCR products were analyzed by agarose gel electrophoresis.

### Small RNA blot assays

Total RNA (20 μg) was separated in 17% polyacrylamide gels containing 0.59 Tris/Borate EDTA and 7 M urea and transferred to positively charged nylon membrane. Synthesis of DNA or LNA oligonucleotide probes labeled with second-generation DIG Oligonucleotide 3’-End Labeling Kit (Roche) and membrane hybridization at 38ºC were done as described (Tomassi *et al*., 2017). An ImageQuant 800 CCD imager (Cytiva) was used to produce digital images from membranes, and band quantification was done using ImageQuant TL version 10.2 (Cytiva). Oligonucleotides used as probes for sRNA blots are listed in Table S1.

### Small RNA sequencing and data analysis

Total RNA quantity, purity and integrity was assessed using a 2100 Bioanalyzer (RNA 6000 Nano kit, Agilent) before submission to BGI (Hong Kong, China) for sRNA library preparation and SE50 high-throughput sequencing on a DNBSEQ-G-400 sequencer. Quality-trimmed reads with adaptor removed, as received from BGI, were processed using the fastx_collapser toolkit (http://hannonlab.cshl.edu/fastx toolkit) (Hannon, 2010) to merge identical sequences while retaining read counts. Each unique, cleaned read was mapped to the forward strand of the corresponding syn-tasiRNA precursor expressed in each sample (Data S3) using a custom Python script, ensuring exact matches with no gasps or mismatches. The script also calculated read counts and RPMs (reads per million mapped reads) for each mapped position.

To evaluate the processing accuracy of syn-tasiRNA precursors, we quantified the proportion of 19– 24 nt sRNA (+) reads mapping within ± 4 nt of the 5′ end of the syn-tasiRNA guide, as previously described (Carbonell *et al*., 2015; Cuperus *et al*., 2010). Additionally, phasing register tables were generated by determining the percentage of 21-nt sRNA (+) reads in each register relative to the predicted syn-tasiRNA cleavage site, considering all 21-nt positions downstream of the cleavage site, following established methods (Carbonell *et al*., 2014; Cisneros *et al*., 2025).

### 5’-RLM-RACE

RNA ligase-mediated rapid amplification of 5′ cDNA ends (5′-RLM RACE) was performed using the GeneRacer kit (Life Technologies), essentially as described (Carbonell *et al*., 2015). Specifically, first-round PCR amplification was done with the GeneRacer 5′ and gene-specific AC-1251 oligonucleotides, followed by a second round PCR using the GeneRacer 5’ nested and AC-1252 oligonucleotides. The resulting 5′-RLM-RACE products were gel-purified, cloned using the Zero Blunt TOPO PCR Cloning Kit (Life Technologies), transformed into *Escherichia coli* DH5α and screened for inserts before sequencing. Control PCR reactions to amplify *SlSFT* and *SlACT* were done using oligonucleotide pairs AC-1250/AC-1252 and AC-280/AC-281, respectively. A complete list of the oligonucleotides used is provided in Table S1.

### Protein blot analysis

Protein separation, membrane transfer, antibody incubation and target protein detection were performed as previously described (Cisneros *et al*., 2025). To assess overall protein content, membranes were stained with Ponceau red S solution (Thermo Fisher Scientific).

### Statistical analysis

The statistical analyses used are detailed in the figure legends. Significant differences were assessed using a two-tailed Student’s *t*-test.

### Gene and virus identifiers

The gene identifiers for *N. benthamiana* and *S. lycopersicum* are as follows: *NbSu* (Nbv5.1tr6204879), *SlACT* (Solyc04g011500.3.1), *SlDXS1* (Solyc01g067890), *SlSFT* (Solyc03g063100.2.1), *SlSu* (Solyc10g008740) and *SlTIP41* (SGN-U584254). The genome identifiers for the TSWV LL-N.05 segment L, M and S are KP008128, FM163373 and KP008129, respectively. PVX-based constructs include PVX sequence variant MT799816.1. *Escherichia coli* β-glucuronidase gene sequence corresponds to GenBank accession number S69414.1.

## ACKNOWLEDGMENTS

We thank Javier Forment (IBMCP) for his support with sRNA sequencing data analysis, Alejandro Atarés (IBMCP) for sharing greenhouse space, and the IBMCP greenhouse staff for their assistance in plant maintenance. This work was supported by grants or fellowships from MCIN/AEI/10.13039/501100011033 and/or by the “European Union NextGenerationEU/PRTR” [PID2021-122186OB-100, PDC2022-133241-I00 and CNS2022135107 to A.C.; PRE2019-088439 and PRE2022103177 to A.E.C. and M.J.M, respectively], and from the European Union’s Horizon Europe research and innovation programme under the Marie Skłodowska-Curie grant agreement No 101149036 (HORIZON-MSCA-2023-PF-01-101149036).

## AUTHOR CONTRIBUTIONS

AHT. and MJ-M did most of the experimental work with the help of AEC, AA, FO and ST-F. SP, TG and AA generated and maintained the transgenic tomato lines. AHT, MJ-M, AEC and AC analyzed the data. A.C. conceived the research, supervised the project and wrote the manuscript with input from the rest of authors.

## DATA AVAILABILITY STATEMENT

All original data will be made available upon request. High-throughput sequencing data can be found in the Sequence Read Archive (SRA) database under accession number PRJNA1220650. New B/c and vectors are available from Addgene: *pENTR-SlmiR482bTS-B/c* (Addgene plasmid #234368, https://www.addgene.org/234368), *pMDC32B-SlmiR482bTS-B/c* (Addgene plasmid #234369, https://www.addgene.org/234369).

## SUPPORTING INFORMATION

**Data S1**. 21-nt siRNAs from *SlSFT* and *SlLRR1*.

**Data S2**. P-SAMS designs of art-sRNA sequences.

**Data S3**. sRNA reads from syn-tasiRNA-expressing tissues.

**Figure S1.** Direct syn-tasiRNA cloning downstream the SlmiR482b target site (TS) in B/c (*Bsa*I/*ccd*B)-based vectors including a *ccd*B cassette flanked by two *Bsa*I sites.

**Figure S2.** Phasing analysis of 21-nt reads corresponding to *SlSFT* and *SlSFT*.

**Figure S3.** Analysis of SlmiR482b and SlmiR6020 presence in *Solanum lycopersicum* agroinfiltrated tissues.

**Table S1**. Name, sequence and use of oligonucleotides used in this study.

**Text S1**. Protocol to design and clone syn-tasiRNAs downstream the 3’D1[+] position in *Bs*aI/*ccd*B-based (‘B/c’) vectors *pENTR-SlmiR482bTS-B/c* and *pMDC32B-SlmiR482bTS-B/c*.

**Text S2**. Protocol to generate PVX-based syn-tasiRNA constructs.

**Text S3**. DNA sequence in FASTA format of all precursors used to express art-sRNAs in plants.

**Text S4**. DNA sequence of *Bsa*I-*ccd*B-based (B/c) vectors used for direct cloning of syn-tasiRNAs.

