## Supplemental Figures-Tables-Texts for "Precision RNAi in Tomato Using Synthetic Trans-Acting Small Interfering RNAs Derived From Minimal Precursors"

### SUPPORTING INFORMATION

**Data S1**. 21-nt siRNAs from *SlSFT* and *SlLRR1*.

**Data S2**. P-SAMS designs of art-sRNA sequences.

**Data S3**. sRNA reads from syn-tasiRNA-expressing tissues.

**Figure S1**. Direct syn-tasiRNA cloning downstream the SlmiR482b target site (TS) in B/c (*Bsa*I/*ccd*B)-based vectors including a *ccd*B cassette flanked by two *Bsa*I sites.

**Figure S2**. Phasing analysis of 21-nt reads corresponding to *SlSFT* and *SlSFT*.

**Figure S3**. Analysis of SlmiR482b and SlmiR6020 presence in *Solanum lycopersicum* agroinfiltrated tissues.

**Table S1**. Name, sequence and use of oligonucleotides used in this study.

**Text S1**. Protocol to design and clone syn-tasiRNAs downstream the 3'D1[+] position in *Bs*aI/*ccd*B-based ('B/c') vectors *pENTR-SlmiR482bTS-B/c* and *pMDC32B-SlmiR482bTS-B/c*.

**Text S2**. Protocol to generate PVX-based syn-tasiRNA constructs.

**Text S3**. DNA sequence in FASTA format of all precursors used to express art-sRNAs in plants.

**Text S4**. DNA sequence of *Bsa*I-*ccd*B-based (B/c) vectors used for direct cloning of syn-tasiRNAs.

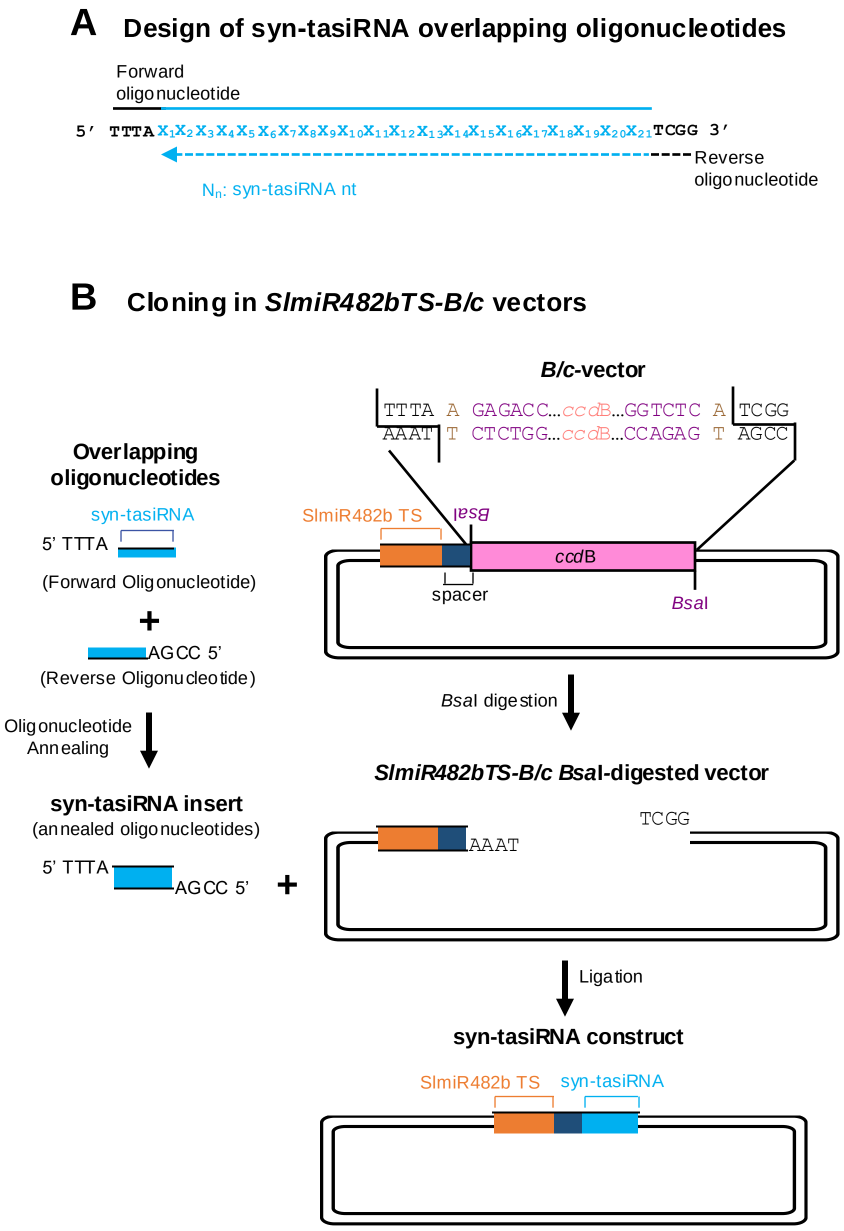


**Figure S1**. Direct syn-tasiRNA cloning downstream the SlmiR482b target site (TS) in *B/c* (*Bsa*I/*ccd*B)-based vectors including a *ccd*B cassette flanked by two *Bsa*I sites. (**A**) Design of two overlapping oligonucleotides for syn-tasiRNA cloning. Sequence covered by the forward and reverse oligonucleotides are represented with continuous or dotted lines, respectively. Nucleotides of the syn-tasiRNA sequence are in blue, and oligonucleotide 5' overhangs are in black and bold. (**B**) Diagram of the steps for syn-tasiRNA cloning in *SlmiR482bTS-B/c* vectors. The syn-tasiRNA insert obtained after annealing the two overlapping oligonucleotides has 5' TTTA and 5'-CCGA overhangs and is directly inserted into the *Bsa*I-linearized *SlmiR482bTS-B/c*-based vectors. Nucleotides of the *Bsa*I sites and arbitrary nucleotides used as spacers between the *Bsa*I recognition site and the *AtTAS1c* sequence are in purple and light brown, respectively. Nucleotides of the SlmiR482b TS are in orange. The *AtTAS1c*-derived spacer and the syn-tasiRNA sequences are in dark and light blue, respectively.

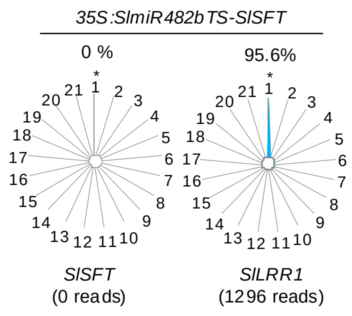


**Figure S2**. Phasing analysis of amiRNA target *SlSFT* mRNA-derived 21 nucleotide small RNAs. Radar plots show proportions of 21-nucleotide reads corresponding to each of the 21 registers from *SlFT*, with position 1 designated as immediately after the amiRNA guided cleavage site. Control plot for tasiRNA-generating *SlLRR1* is shown. The percentage of 21-nucleotide reads corresponding to phasing register 1 is indicated.

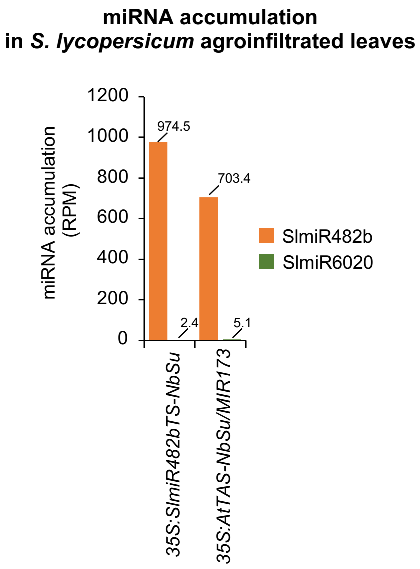


**Figure S3**. Analysis of SlmiR482b and SlmiR6020 presence in *Solanum lycopersicum* agroinfiltrated tissues. Left, bar graph showing the accumulation (reads per million, RPM) of SlmiR482b and SlmiR6020 revealed by high-throughput sequencing of small RNA libraries prepared from leaves agroinfiltrated with different constructs.

**Table S1.** Name, sequence and use of oligonucleotides used in this study.

| Name | Sequence | Type* | Construct/Aim |
|---|---|---|---|
| AC-50 | CCGATTCACCCAGAGCTGATA | ssDNA | Probe to detect syn-tasiR-TSWV-2 |
| AC-51 | CGGTATGCCCCACTATACCAA | ssDNA | Probe to detect syn-tasiR-TSWV-3 |
| AC-52 | AAGATGGATTGTGCACTCTGA | ssDNA | Probe to detect syn-tasiR-TSWV-4 |
| AC-55 | AGGGGCCATGCTAATCTTCTC | ssDNA | Probe for U6 detection |
| AC-280 | CTAGGCTGGGTTCGCAGGAGATGATGC | ssDNA | *SlACT* qPCR |
| AC-281 | GTCTTTTTGACCCATACCCACCATCACAC | ssDNA | |
| AC-416 | A+GGA+CAC+AAT+CAC+GTC+TTA+CA | ssLNA | Probe to detect syn-tasiR-TSWV-1 |
| AC-417 | G+CGG+GAA+GTC+CAC+CAC+GGT+TA | ssLNA | Probe for syn-tasiR-NbSu detection |
| AC-452 | ATGGAGTTTTTGAGTCTTCTGC | ssDNA | *SlTIP41* qPCR |
| AC-453 | GCTGCGTTTCTGGCTTAGG | ssDNA | |
| AC-525 | | | |
| AC-654 | GGGAATCAATCACAGTGTTGGC | ssDNA | syn-tasiRNA precursors detection |
| AC-655 | GCTACTATGGCACGGGCTGTAC | ssDNA | |
| AC-657 | ATGTCAGGCCTGTTCACTATCC | ssDNA | PVX detection |
| AC-658 | TGGTGGTGGTAGAGTGACAAC | ssDNA | |
| AC-775 | agaggtcagcaccagctagcGGCATGGGCGGTGTAGG CAAGATAGACCATTTATCCCCCTTGTATGAT TCCCGCTCTCAATGTCGATTGTCCCAGagggttt gttaagtttccct | dsDNA | *35S:PVX-SlmiR482bTS-SlSu(x2)* |
| AC-776 | agaggtcagcaccagctagcGGCATGGGCGGTGTAGG CAAGATAGACCATTTATATAGGACTGTGTC TTGGCAGTATTGATCCTTCTTCGACCGAaggg tttgttaagtttccct | dsDNA | *35S:PVX-SlmiR482bTS-SlDXS(x2)* |
| AC-820 | TGACCATGGATCTCCTGTTG | ssDNA | *SlDXS* qPCR |
| AC-821 | GCCTCTCTGGTTTGTCCAAG | ssDNA | |
| AC-822 | CTACACCCTTCACTTCTCGTTCC | ssDNA | *SlSu* qPCR |
| AC-823 | AGTTTCTGAGCCTGTTCTTGAGC | ssDNA | |
| AC-824 | TGTAGGCATGGGCGGTGTAGGCAAGATAGA CCATTTATATTGACCCACACTTTGCCGA | ssDNA | *35S:SlmiR482bTS-GUS*$_{Sl}$ |
| AC-825 | AATGTCGGCAAAGTGTGGGTCAATATAAAT GGTCTATCTTGCCTACACCGCCCATGCC | ssDNA | |
| AC-826 | TGTAGGCATGGGCGGTGTAGGCAAGATAGA CCATTTATGTATGACTCCCGGAATTCCA | ssDNA | *35S:SlmiR482bTS-NbSu* |
| AC-827 | AATGTGGAATTCCGGGAGTCATACATAAAT GGTCTATCTTGCCTACACCGCCCATGCC | ssDNA | |
| AC-828 | TGTAAAAGATACTCGGAAAACATTTATAGA CCATTTATATTGACCCACACTTTGCCGA | ssDNA | *35S:SlmiR6020TS-GUS*$_{Sl}$ |
| AC-829 | AATGTCGGCAAAGTGTGGGTCAATATAAAT GGTCTATAAATGTTTTCCGAGTATCTTT | ssDNA | |
| AC-830 | TGTAAAAGATACTCGGAAAACATTTATAGA CCATTTATGTATGACTCCCGGAATTCCA | ssDNA | *35S:SlmiR6020TS-NbSu* |
| AC-831 | AATGTGGAATTCCGGGAGTCATACATAAAT GGTCTATAAATGTTTTCCGAGTATCTTT | ssDNA | |
| AC-832 | GTTGGTCGTGTGGTAGGGGA | ssDNA | *SlSFT* qPCR |
| AC-833 | GCCTAAGCTCGCATCCATTA | ssDNA | |
| AC-874 | G+CGG+GAA+TCA+TAC+AAG+GGG+GA | ssLNA | Probe to detect syn-tasiR-SlSu-1 |
| AC-875 | C+TGG+GAC+AAT+CGA+CAT+TGA+GA | ssLNA | Probe to detect syn-tasiR-SlSu-2 |
| AC-876 | C+TGC+CAA+GAC+ACA+GTC+CTA+TA | ssLNA | Probe to detect syn-tasiR-SlDXS-1 |
| AC-877 | T+CGG+TCG+AAG+AAG+GAT+CAA+TA | ssLNA | Probe to detect syn-tasiR-SlDXS-2 |
| AC-892 | TGTAGGCATGGGCGGTGTAGGCAAGATAGA CCATTTATCCCCCTTGTATGATTCCCGCTCT CAATGTCGATTGTCCCAG | ssDNA | *35S:SlmiR482bTS-SlSu(x2)* |
| AC-893 | AATGCTGGGACAATCGACATTGAGAGCGGG AATCATACAAGGGGGATAAATGGTCTATCT TGCCTACACCGCCCATGCC | ssDNA | |
| AC-894 | TGTAGGCATGGGCGGTGTAGGCAAGATAGA CCATTTATATAGGACTGTGTCTTGGCAGTAT TGATCCTTCTTCGACCGA | ssDNA | *35S:SlmiR482bTS-SlDXS(x2)* |
| AC-895 | AATGTCGGTCGAAGAAGGATCAATACTGCC AAGACACAGTCCTATATAAATGGTCTATCT TGCCTACACCGCCCATGCC | ssDNA | |

| | | | |
|---|---|---|---|
| AC-902 | TGTAGGCATGGGCGGTGTAGGCAAGATAGA CCATTTAaGAGACCggtctcATCGG | ssDNA | *pENTR-SlmiR482bTS-BB, pMDC32B-SlmiR482bTS-BB* |
| AC-903 | AATGCCGATgagaccGGTCTCtTAAATGGTCTA TCTTGCCTACACCGCCCATGCC | ssDNA | |
| AC-909 | TTTATGCATACACTGTTTGCCGGCT | ssDNA | *35S:SlmiR482bTS-SlSFT* |
| AC-910 | CCGAAGCCGGCAAACAGTGTATGCA | ssDNA | |
| AC-915 | TGTAGGCATGGGCGGTGTAGGCAAGATAGA CCATTTATATTGACCCACACTTTGCCGATAA CCTTCACCCGGTTGCCAC | ssDNA | *35S:SlmiR482bTS-GUS$_{Sl}$(x2)* |
| AC-916 | AATGGTGGCAACCGGGTGAAGGTTATCGGC AAAGTGTGGGTCAATATAAATGGTCTATCT TGCCTACACCGCCCATGCC | ssDNA | |
| AC-993 | agaggtcagcaccagctagcGGCATGGGCGGTGTAGG CAAGATAGACCATTTATATTGACCCACACT TTGCCGATAACCTTCACCCGGTTGCCACTAT TGACCCACACTTTGCCGATAACCTTCACCC GGTTGCCACagggtttgttaagtttccct | dsDNA | *35S:PVX-SlmiR482bTS-GUS$_{Sl}$(x4)* |
| AC-1205 | agaggtcagcaccagctagcGTGATTTTTCTCTACAAG CGAATAGACCATTTATGTAAGACGTGATTG TGTCCTTATCAGCTCTGGGTGAATCGGTTGG TATAGTGGGGCATACCGTCAGAGTGCACAA TCCATCTTagggtttgttaagtttccct | dsDNA | *35S:PVX-AtmiR173aTS-TSWV(x4)* |
| AC-1224 | A+GCC+GGC+AAA+CAG+TGT+ATG+CA | LNA | Probe to detect syn-tasiR-SlSFT |
| AC-1250 | AGGGTTGAAGTTGGAGGAGATGACC | ssDNA | *SlSFT* control PCR (5'RLM-RACE) |
| AC-1251 | GTAAGAGAAGTAGTAGATATTGGTGGTT | ssDNA | *SlSFT* round 1 PCR (5'RLM-RACE) |
| AC-1252 | CGTCCACCACTGCCACTCTCTCTTTG | ssDNA | *SlSFT* control PCR (5'RLM-RACE)<br>*SlSFT* round 2 PCR (5'RLM-RACE) |
| GeneRacer 5' Oligo | CGACTGGAGCACGAGGACACTGA | ssDNA | *SlSFT* round 1 PCR (5'RLM-RACE) |
| GeneRacer 5' Nested Oligo | GGACACTGACATGGACTGAAGGAGTA | ssDNA | *SlSFT* round 2 PCR (5'RLM-RACE) |
| GeneRacer Oligo dT | GCTGTCAACGATACGCTACGTAACGGCATG ACAGTGTTTTTTTTTTTTTTTTTTTTTTTT | ssDNA | cDNA synthesis (5'RLM-RACE) |
| GeneRacer RNA Oligo Adapter | CGACUGGAGCACGAGGACACUGACAUGGA CUGAAGGAGUAGAAA | ssRNA | RNA ligation (5'RLM-RACE) |

*ssDNA: single-stranded DNA; dsDNA: double-stranded DNA; LNA: locked nucleic acid;ssRNA, single-stranded RNA.

**Text S1.** Protocol to design and clone syn-tasiRNAs downstream the 3'D1[+] position in *Bsa*I/*ccd*B-based ('B/c') vectors *pENTR-SlmiR482bTS-B/c* and *pMDC32B-SlmiR482bTS-B/c*.

**1. Selection of the syn-tasiRNA sequence(s)**

Use the Syn-tasiRNA Designer app from the P-SAMS webtool at http://p-sams.carringtonlab.org/syntasi/designer.

**2. Design of syn-tasiRNA oligonucleotides for cloning**

Next are described some designs for cloning two syn-tasiRNAs in tandem downstream the 3'D1[+] position.

Use vectors *pENTR-SlmiR482bTS-B/c* or *pMDC32-SlmiR482bTS-B/c* and order the following oligos:

-Forward oligonucleotide (46 b):

**TTTA**$X_1X_2X_3X_4X_5X_6X_7X_8X_9X_{10}X_{11}X_{12}X_{13}X_{14}X_{15}X_{16}X_{17}X_{18}X_{19}X_{20}X_{21}X_1X_2X_3X_4X_5X_6X_7X_8X_9X_{10}X_{11}X_{12}$
$X_{13}X_{14}X_{15}X_{16}X_{17}X_{18}X_{19}X_{20}X_{21}$

-Reverse oligonucleotide (46 b):

**CCGA**$Y_{21}Y_{20}Y_{19}Y_{18}Y_{17}Y_{16}Y_{15}Y_{14}Y_{13}Y_{12}Y_{11}Y_{10}Y_9Y_8Y_7Y_6Y_5Y_4Y_3Y_2Y_1Y_{21}Y_{20}Y_{19}Y_{18}Y_{17}$
$Y_{16}Y_{15}Y_{14}Y_{13}Y_{12}Y_{11}Y_{10}Y_9Y_8Y_7Y_6Y_5Y_4Y_3Y_2Y_1$

Where:

$X_1X_2X_3X_4X_5X_6X_7X_8X_9X_{10}X_{11}X_{12}X_{13}X_{14}X_{15}X_{16}X_{17}X_{18}X_{19}X_{20}X_{21}$=syn-tasiRNA-1 sequence

$X_1X_2X_3X_4X_5X_6X_7X_8X_9X_{10}X_{11}X_{12}X_{13}X_{14}X_{15}X_{16}X_{17}X_{18}X_{19}X_{20}X_{21}$=syn-tasiRNA-2 sequence

$Y_{21}Y_{20}Y_{19}Y_{18}Y_{17}Y_{16}Y_{15}Y_{14}Y_{13}Y_{12}Y_{11}Y_{10}Y_9Y_8Y_7Y_6Y_5Y_4Y_3Y_2Y_1$=syn-tasiRNA-1 reverse-complement sequence

$Y_{21}Y_{20}Y_{19}Y_{18}Y_{17}Y_{16}Y_{15}Y_{14}Y_{13}Y_{12}Y_{11}Y_{10}Y_9Y_8Y_7Y_6Y_5Y_4Y_3Y_2Y_1$=syn-tasiRNA-2 reverse-complement sequence

**Example**

The sequences of the two oligonucleotides to clone syn-tasiRNAs 'syn-tasiR-TRY' (TCCCATTCGATACTGCTCGCC) and 'syn-tasiR-Ft' (TTGGTTATAAAGGAAGAGGCC) in positions 3'D2[+] and 3'D3[+], respectively, of minimal precursors included in *SlmiR482bTS*-based B/c vectors are:

-Forward oligonucleotide (46 b):

**TTTA**TCCCATTCGATACTGCTCGCCTTGGTTATAAAGGAAGAGGCC

-Reverse oligonucleotide (46 b):

**CCGA**GGCCTCTTCCTTTATAACCAAGGCGAGCAGTATCGAATGGGA

### 3. Cloning of the syn-tasiRNA sequence(s) in B/c-based vectors

*Notes:*

*-New available -B/c vectors are listed in Table I at the end of the section.*

-B/c-*based vectors must be propagated in a* ccd*B resistant* E. coli *strain such as DB3.1.*

-Alternatively, Bsa*I digestion of the B/c vector and subsequent ligation of the amiRNA oligonucleotide insert can be done in separate reactions*

#### 3.1. Oligonucleotide annealing

-Dilute sense oligonucleotide and antisense oligonucleotide in sterile $H_2O$ to a final concentration of 100 µM.

-Prepare Oligo Annealing Buffer:

60 mM Tris-HCl (pH 7.5)

500 mM NaCl

60 mM $MgCl_2$

10 mM DTT

***Note:*** *Prepare 1 ml aliquots of Oligo Annealing Buffer and store at -20$^0$C.*

-Assemble the annealing reaction in a PCR tube as described below:

| | |
|---|---|
| Forward oligonucleotide (100 µM) | 2 µL |
| Reverse oligonucleotide (100 µM) | 2 µL |
| Oligo Annealing Buffer | 46 µL |
| Total volume | 50 µL |

The final concentration of each oligonucleotide is 4 µM.

-Use a thermocycler to heat the annealing reaction 5 min at 94°C and then cool down (0.05ºC/sec) to 20°C.

-Dilute the annealed oligonucleotides just prior to assembling the digestion-ligation reaction as described below:

| | |
|---|---|
| Annealed oligonucleotides | 3 µL |
| $dH_2O$ | 37 µL |
| Total volume | 40 µL |

The final concentration of each oligonucleotide is 0.15 µM.

***Note**: Do not store the diluted oligonucleotides.*

3.2. Digestion-ligation reaction

- Assemble the digestion-ligation reaction as described below:

| | |
|---|---|
| B/c vector (x ug/uL) | Y µL (50 ng) |
| Diluted annealed oligonucleotides | 1 µL |
| 10x T4 DNA ligase buffer | 1 µL |
| T4 DNA ligase (400 U/µL) | 1 µL |
| *Bsa*I (10U/ µL, NEB) | 1 µL |
| $dH_2O$ | to 10 µL |
| Total volume | 10 µL |

Prepare a negative control reaction lacking *Bsa*I.

-Mix the reactions by pipetting. Incubate the reactions at room temperature for 5 minutes at 37°C.

3.3. *E.coli* transformation and analysis of transformants

-Transform 1-5 ul of the digestion-ligation reaction into an *E. coli* strain that doesn't have *ccd*B resistance (e.g. DH10B, TOP10, …) to do counter-selection.

-Pick two colonies/construct, grow LB-Kan (100 mg/ml) cultures and purify plasmids.

-Sequence with appropriate primers: M13-F (CCCAGTCACGACGTTGTAAAACGACGG) and M13-R (CAGAGCTGCCAGGAAACAGCTATGACC) for *pENTR*-based vectors; attB1 (ACAAGTTTGTACAAAAAAGCAGGCT) and attB2 (ACCACTTTGTACAAGAAAGCTGGGT) primers for *pMDC32B*-based vectors).

**Table I**: *Bsa*I/*ccd*B-based ('B/c') vectors for direct cloning of syn-tasiRNAs downstream position 3'D1[+] in minimal precursor including SlmiR482b TS.

| Vector | Small RNA expressed | Bacterial antibiotic resistance | Plant antibiotic resistance | GATEWAY use | Backbone | Promoter of syn-tasiRNA cassette | Terminator of syn-tasiRNA cassette | Plant species tested |
|---|---|---|---|---|---|---|---|---|
| *pENTR-SlmiR482bTS-B/c* | syn-tasiRNAs | Kanamycin | – | Donor | *pENTR* | – | – | – |
| *pMDC32B-SlmiR482bTS-B/c* | syn-tasiRNAs | Kanamycin Hygromycin | Hygromycin | – | *pMDC32* | *CaMV* 2x35S | *Nos* | *S. lycopersicum* |

**Text S2.** Protocol to generate PVX-based syn-tasiRNA constructs.

### 1. Preparation of the dsDNA syn-tasiRNA insert

Design and order a dsDNA (eg. ultramer duplex in IDT) including the sequences of your syn-tasiRNA(s) (2 in the following example) following the 22-nt miRNA target site of interest, as follows:

agaggtcagcaccagctagc$X_1X_2X_3X_4X_5X_6X_7X_8X_9X_{10}X_{11}X_{12}X_{13}X_{14}X_{15}X_{16}X_{17}X_{18}X_{19}X_{20}X_{21}X_{22}$TAGACCATTTA$X_1X_2X_3X_4X_5X_6X_7X_8X_9X_{10}X_{11}X_{12}X_{13}X_{14}X_{15}X_{16}X_{17}X_{18}X_{19}X_{20}X_{21}$$X_1X_2X_3X_4X_5X_6X_7X_8X_9X_{10}X_{11}X_{12}X_{13}X_{14}X_{15}X_{16}X_{17}X_{18}X_{19}X_{20}X_{21}$agggtttgttaagtttccct

Where:

- -X is a DNA base of the 22-nt miRNA target site sequence, and the subscript number is the base position
- -X is a DNA base of the syn-tasiRNA-1 sequence, and the subscript number is the base position in the syn-tasiRNA* 21-mer
- -X is a DNA base of the syn-tasiRNA-2 sequence, and the subscript number is the base position in the syn-tasiRNA 21-mer
- -x is a DNA base of the PVX sequence, required for Gibson-based assembly
- -X is a DNA base of the *AtTAS1c* sequence

Note that:

- -In general, **$X_1$=T** and **$X_1$=T** for amiRNA association with AGO1.

Fragment #1 (syn-tasiRNA precursor) is ready.

### 2. Preparation of the vector

-Digest *pLB-PVX* with *Mlu*I.

-Gel purify the 9921 bp band corresonding to linearized plasmid.

-Quantify 1 ul in Nanodrop.

Fragment #2 (backbone vector) is ready.

### 3. Assembly

-Assemble the Gibbson reaction as described below:

| | |
|---|---|
| Fragment 1 (dsDNA insert)[a] | |
| Fragment 2 (vector)[b,c,d] | |
| GeneArt Gibson Assembly HiFI Master Mix | 5 µL |
| $dH_2O$ | to 10 µL |

Total volume 10 µL

[a]The optimal amount of vector is between 50-100 ng

[b]Insert/vector molar excess is between 2-3.

[c]Total DNA amount is between 0.02-0.5 pmol

[d]Mass to moles conversions can be calculated here:
http://nebiocalculator.neb.com/#!/ssdnaamt

-Incubate reactions at 50ºC for 1h.

-Clean up reactions with a column (e.g. Zymo Research)

-Transform 1-4 µL in *E. coli* DH5α

-Plate in L-Kan plates and incubate 16h at 37ºC

### 4. Clone verification

-Pick several colonies and grow in liquid LB-Kan 16h at 37ºC, and purify plasmids.

-Digest candidate clones with *Apa*I+*Xho*I

Good clones: 8595 + **1409** bp

Bad clones (empty *pLB-PVX*): 9921 bp + **1738** bp

-Confirm insert sequence by Sanger sequencing with forward and reverse oligos AC-654 (`GGGAATCAATCACAGTGTTGGC`) and/or AC-655 (`GCTACTATGGCACGGGCTGTAC`), respectively.

**Text S3**. DNA sequence in FASTA format of all precursors used to express syn-tasiRNAs in plants.

### 1. *AtTAS1c*-based precursors

***>AtTAS1c-NbSu***

AAACCTAAACCTAAACGGCTAAGCCCGACGTCAAATACCAAAAAGAGAAAAACAAGAGCGCCGTCAAGCTCTGCAAATACGATCTGTAAG
TCCATCTTAACACAAAAGTGAGATGGGTTCTTAGATCATGTTCCGCCGTTAGATCGAGTCATGGTCTTGTCTCATAGAAAGGTACTTTCG
TTTACTTCTTTTGAGTATCGAGTAGAGCGTCGTCTATAGTTAGTTTGAGATTGCGTTTGTCAGAAGTTAGGTTCAATGTCCCGGTCCAAT
TTTCACCAGCCATGTGTCAGTTTCGTTCCTTCCCGTCCTCTTCTTTGATTTCGTTGGGTTACGGATGTTTTCGAGATGAAACAGCATTGT
TTTGTTGTGATTTTTCTCTACAAGCGAATAGACCATTTATGTATGACTCCCGGAATTCCATCGGTGGATCTTAGAAAATTATTCTAAGTC
CAACATAGCGTATTCTAAGTTCAACATATCGACGAACTAGAAAAGACATTGGACATATTCCAGGATATGCAAAAGAAAACAATGAATATT
GTTTTGAATGTGTTCAAGTAAATGAGATTTTCAAGTCGTCTAAAGAACAGTTGCTAATACAGTTACTTATTTCAATAAATAATTGGTTCT
AATAATACAAAACATATTCGAGGATATGCAGAAAAAAAGATGTTTGTTATTTTGAAAAGCTTGAGTAGTTTCTCTCCGAGGTGTAGCGAA
GAAGCATCATCTACTTTGTAATGTAATTTTCTTTATGTTTTCACTTTGTAATTTTATTTGTGTTAATGTACCATGGCCGATATCGGTTTT
ATTGAAAGAAAATTTATGTTACTTCTGTTTTGGCTTTGCAATCAGTTATGCTAGTTTTCTTATACCCTTTCGTAAGCTTCCTAAGGAATC
GTTCATTGATTTCCACTGCTTCATTGTATATTAAAACTTTACAACTGTATCGACCATCATATAATTCTGGGTCAAGAGATGAAAATAGAA
CACCACATCGTAAAGTGAAAT

*AtTAS1c*
AtmiR173a TS
syn-tasiR-NbSu

### 2. Minimal syn-tasiRNA precursors

***>SlmiR482bTS-GUS$_{Nb}$***

GGCATGGGCGGTGTAGGCAAGATAGACCATTTATATTGACCCACACTTTGCCGA

*AtTAS1c*
SlmiR482b TS
syn-tasiR-GUS$_{Nb}$

***>SlmiR482bTS-NbSu***

GGCATGGGCGGTGTAGGCAAGATAGACCATTTATGTATGACTCCCGGAATTCCA

*AtTAS1c*
SlmiR482b TS
syn-tasiR-NbSu

***>SlmiR6020TS-GUS$_{Nb}$***

AAAGATACTCGGAAAACATTTATAGACCATTTATATTGACCCACACTTTGCCGA

*AtTAS1c*
SlmiR6020 TS
syn-tasiR-GUS$_{Nb}$

***>SlmiR6020TS-NbSu***

AAAGATACTCGGAAAACATTTATAGACCATTTATGTATGACTCCCGGAATTCCA

*AtTAS1c*
SlmiR6020 TS
syn-tasiR-NbSu

***>SlmiR482bTS-SlSFT***

GGCATGGGCGGTGTAGGCAAGATAGACCATTTATGCATACACTGTTTGCCGGCT

*AtTAS1c*
SlmiR482b TS
syn-tasiR-SlSFT

***>SlmiR482bTS-GUS$_{S1}$(x4)***

GGCATGGGCGGTGTAGGCAAGATAGACCATTTATATTGACCCACACTTTGCCGATAACCTTCACCCGGTTGCCACTATTGACCC
ACACTTTGCCGATAACCTTCACCCGGTTGCCAC

*AtTAS1c*
SlmiR482b TS
syn-tasiR-GUS$_{S1}$
syn-tasiR-GUS$_{S1}$-2

***>SlmiR482bTS-TSWV(x4)***

GGCATGGGCGGTGTAGGCAAGATAGACCATTTATGTAAGACGTGATTGTGTCCTTATCAGCTCTGGGTGAATCGGTTGGTATAG
TGGGGCATACCGTCAGAGTGCACAATCCATCTT

*AtTAS1c*
SlmiR482b TS
syn-tasiR-TSWV-1
syn-tasiR-TSWV-2
syn-tasiR-TSWV-3
syn-tasiR-TSWV-4

***>AtmiR173a-TSWV(x4)***

GTGATTTTTCTCTACAAGCGAATAGACCATTTATGTAAGACGTGATTGTGTCCTTATCAGCTCTGGGTGAATCGGTTGGTATAG
TGGGGCATACCGTCAGAGTGCACAATCCATCTT

*AtTAS1c*
AtmiR173a TS
syn-tasiR-TSWV-1
syn-tasiR-TSWV-2
syn-tasiR-TSWV-3
syn-tasiR-TSWV-4

***>SlmiR482bTS-SlGUS$_{S1}$(x2)***

GGCATGGGCGGTGTAGGCAAGATAGACCATTTATATTGACCCACACTTTGCCGATAACCTTCACCCGGTTGCCAC

*AtTAS1c*
SlmiR482b TS
syn-tasiR-GUS$_{S1}$-1
syn-tasiR-GUS$_{S1}$-2

***>SlmiR482bTS-SlSu(x2)***

GGCATGGGCGGTGTAGGCAAGATAGACCATTTATCCCCCTTGTATGATTCCCGCTCTCAATGTCGATTGTCCCAG

*AtTAS1c*
SlmiR482b TS
syn-tasiR-SlSu-1
syn-tasiR-SlSu-2

***>SlmiR482bTS-SlDXS(x2)***

GGCATGGGCGGTGTAGGCAAGATAGACCATTTATATAGGACTGTGTCTTGGCAGTATTGATCCTTCTTCGACCGA

*AtTAS1c*
SlmiR482b TS
syn-tasiR-SlDXS1-1
syn-tasiR-SlDXS1-2

**Text S4.** DNA sequence of *Bsa*I-*ccd*B-based (B/c) vectors used for direct cloning of syn-tasiRNAs.

***>pENTR-SlmiR482bTS-B/c*** (4082 bp)

CTTTCCTGCGTTATCCCCTGATTCTGTGGATAACCGTATTACCGCCTTTGAGTGAGCTGATACCGCTCGCCGCAGCCGAACGACCGAGCG
CAGCGAGTCAGTGAGCGAGGAAGCGGAAGAGCGCCCAATACGCAAACCGCCTCTCCCCGCGCGTTGGCCGATTCATTAATGCAGCTGGCA
CGACAGGTTTCCCGACTGGAAAGCGGGCAGTGAGCGCAACGCAATTAATACGCGTACCGCTAGCCAGGAAGAGTTTGTAGAAACGCAAAA
AGGCCATCCGTCAGGATGGCCTTCTGCTTAGTTTGATGCCTGGCAGTTTATGGCGGGCGTCCTGCCCGCCACCCTCCGGGCCGTTGCTTC
ACAACGTTCAAATCCGCTCCCGGCGGATTTGTCCTACTCAGGAGAGCGTTCACCGACAAACAACAGATAAAACGAAAGGCCCAGTCTTCC
GACTGAGCCTTTCGTTTTATTTGATGCCTGGCAGTTCCCTACTCTCGCGTTAACGCTAGCATGGATGTTTTCCCAGTCACGACGTTGTAA
AACGACGGCCAGTCTTAAGCTCGGGCCCCAAATAATGATTTTATTTTGACTGATAGTGACCTGTTCGTTGCAACAAATTGATGAGCAATG
CTTTTTTATAATGCCAACTTTGTACAAAAAAGCAGGCTCCGCGGCCGCCCCCTTCACCTGTA**GGCATGGGCGGTGTAGGCAAGA**TAGACC
**ATTTA**AGAGACCATTAGGCACCCCAGGCTTTACACTTTATGCTTCCGGCTCGTATAATGTGTGGATTTTGAGTTAGGAGCCGTCGAGATT
TTCAGGAGCTAAGGAAGCTAAAATGGAGAAAAAAATCACTGGATATACCACCGTTGATATATCCCAATGGCATCGTAAAGAACATTTTGA
GGCATTTCAGTCAGTTGCTCAATGTACCTATAACCAGACCGTTCAGCTGGATATTACGGCCTTTTTAAAGACCGTAAAGAAAAATAAGCA
CAAGTTTTATCCGGCCTTTATTCACATTCTTGCCCGCCTGATGAATGCTCATCCGGAGTTCCGTATGGCAATGAAAGACGGTGAGCTGGT
GATATGGGATAGTGTTCACCCTTGTTACACCGTTTTCCATGAGCAAACTGAAACGTTTTCATCGCTCTGGAGTGAATACCACGACGATTT
CCGGCAGTTTCTACACATATATTCGCAAGATGTGGCGTGTTACGGTGAAAACCTGGCCTATTTCCCTAAAGGGTTTATTGAGAATATGTT
TTTCGTCTCAGCCAATCCCTGGGTGAGTTTCACCAGTTTTGATTTAAACGTGGCCAATATGGACAACTTCTTCGCCCCCGTTTTCACCAT
GGGCAAATATTATACGCAAGGCGACAAGGTGCTGATGCCGCTGGCGATTCAGGTTCATCATGCCGTTTGTGATGGCTTCCATGTCGGCAG
AATGCTTAATGAATTACAACAGTACTGCGATGAGTGGCAGGGCGGGGCGTAAACGCGTGGAGCCGGCTTACTAAAAGCCAGATAACAGTA
TGCGTATTTGCGCGCTGATTTTTGCGGTATAAGAATATATACTGATATGTATACCCGAAGTATGTCAAAAAGAGGTATGCTATGAAGCAG
CGTATTACAGTGACAGTTGACAGCGACAGCTATCAGTTGCTCAAGGCATATATGATGTCAATATCTCCGGTCTGGTAAGCACAACCATGC
AGAATGAAGCCCGTCGTCTGCGTGCCGAACGCTGGAAAGCGGAAAATCAGGAAGGGATGGCTGAGGTCGCCCGGTTTATTGAAATGAACG
GCTCTTTTGCTGACGAGAACAGGGGCTGGTGAAATGCAGTTTAAGGTTTACACCTATAAAAGAGAGAGCCGTTATCGTCTGTTTGTGGAT
GTACAGAGTGATATTATTGACACGCCCGGCCGACGGATGGTGATCCCCCTGGCCAGTGCACGTCTGCTGTCAGATAAAGTCTCCCGTGAA
CTTTACCCGGTGGTGCATATCGGGGATGAAAGCTGGCGCATGATGACCACCGATATGGCCAGTGTGCCGGTTTCCGTTATCGGGGAAGAA
GTGGCTGATCTCAGCCACCGCGAAAATGACATCAAAAACGCCATTAACCTGATGTTCTGGGGAATATAAATGTCAGGCTCCCTTATACAC
AGCCAGTCTGCACCTCGACGGTCTCACATTAAGGGTGGGCGCGCCGACCCAGCTTTCTTGTACAAAGTTGGCATTATAAGAAAGCATTGC
TTATCAATTTGTTGCAACGAACAGGTCACTATCAGTCAAAATAAAATCATTATTTGCCATCCAGCTGATATCCCCTATAGTGAGTCGTAT
TACATGGTCATAGCTGTTTCCTGGCAGCTCTGGCCCGTGTCTCAAAATCTCTGATGTTACATTGCACAAGATAAAAATATATCATCATGA
ACAATAAAACTGTCTGCTTACATAAACAGTAATACAAGGGGTGTTATGAGCCATATTCAACGGGAAACGTCGAGGCCGCGATTAAATTCC
AACATGGATGCTGATTTATATGGGTATAAATGGGCTCGCGATAATGTCGGGCAATCAGGTGCGACAATCTATCGCTTGTATGGGAAGCCC
GATGCGCCAGAGTTGTTTCTGAAACATGGCAAAGGTAGCGTTGCCAATGATGTTACAGATGAGATGGTCAGACTAAACTGGCTGACGGAA
TTTATGCCTCTTCCGACCATCAAGCATTTTATCCGTACTCCTGATGATGCATGGTTACTCACCACTGCGATCCCCGGAAAAACAGCATTC
CAGGTATTAGAAGAATATCCTGATTCAGGTGAAAATATTGTTGATGCGCTGGCAGTGTTCCTGCGCCGGTTGCATTCGATTCCTGTTTGT
AATTGTCCTTTTAACAGCGATCGCGTATTTCGTCTCGCTCAGGCGCAATCACGAATGAATAACGGTTTGGTTGATGCGAGTGATTTTGAT
GACGAGCGTAATGGCTGGCCTGTTGAACAAGTCTGGAAAGAAATGCATAAACTTTTGCCATTCTCACCGGATTCAGTCGTCACTCATGGT
GATTTCTCACTTGATAACCTTATTTTTGACGAGGGGAAATTAATAGGTTGTATTGATGTTGGACGAGTCGGAATCGCAGACCGATACCAG
GATCTTGCCATCCTATGGAACTGCCTCGGTGAGTTTTCTCCTTCATTACAGAAACGGCTTTTTCAAAAATATGGTATTGATAATCCTGAT
ATGAATAAATTGCAGTTTCATTTGATGCTCGATGAGTTTTTCTAATCAGAATTGGTTAATTGGTTGTAACACTGGCAGAGCATTACGCTG
ACTTGACGGGACGGCGCAAGCTCATGACCAAAATCCCTTAACGTGAGTTACGCGTCGTTCCACTGAGCGTCAGACCCCGTAGAAAAGATC
AAAGGATCTTCTTGAGATCCTTTTTTTCTGCGCGTAATCTGCTGCTTGCAAACAAAAAAACCACCGCTACCAGCGGTGGTTTGTTTGCCG
GATCAAGAGCTACCAACTCTTTTTCCGAAGGTAACTGGCTTCAGCAGAGCGCAGATACCAAATACTGTCCTTCTAGTGTAGCCGTAGTTA
GGCCACCACTTCAAGAACTCTGTAGCACCGCCTACATACCTCGCTCTGCTAATCCTGTTACCAGTGGCTGCTGCCAGTGGCGATAAGTCG
TGTCTTACCGGGTTGGACTCAAGACGATAGTTACCGGATAAGGCGCAGCGGTCGGGCTGAACGGGGGGTTCGTGCACACAGCCCAGCTTG
GAGCGAACGACCTACACCGAACTGAGATACCTACAGCGTGAGCATTGAGAAAGCGCCACGCTTCCCGAAGGGAGAAAGGCGGACAGGTAT
CCGGTAAGCGGCAGGGTCGGAACAGGAGAGCGCACGAGGGAGCTTCCAGGGGGAAACGCCTGGTATCTTTATAGTCCTGTCGGGTTTCGC
CACCTCTGACTTGAGCGTCGATTTTTGTGATGCTCGTCAGGGGGGCGGAGCCTATGGAAAAACGCCAGCAACGCGGCCTTTTTACGGTTC
CTGGCCTTTTGCTGGCCTTTTGCTCACATGTT

**SlmiR482b target site**
***AtTAS1c*-derived spacer**
M13-F binding site
M13-Reverse binding site
attL1
attL2
Chloramphenicol resistance gene
*ccd*B gene
*Bsa*I site
inverted *Bsa*I site
Kanamycin resistance gene

***>pMDC32B-SlmiR482bTS-B/c*** (11635 bp)

CCAGCCAGCCAACAGCTCCCCGACCGGCAGCTCGGCACAAAATCACCACTCGATACAGGCAGCCCATCAGTCCGGGACGGCGTCAGCGGG
AGAGCCGTTGTAAGGCGGCAGACTTTGCTCATGTTACCGATGCTATTCGGAAGAACGGCAACTAAGCTGCCGGGTTTGAAACACGGATGA
TCTCGCGGAGGGTAGCATGTTGATTGTAACGATGACAGAGCGTTGCTGCCTGTGATCACCGCGGTTTCAAAATCGGCTCCGTCGATACTA
TGTTATACGCCAACTTTGAAAACAACTTTGAAAAAGCTGTTTTCTGGTATTTAAGGTTTTAGAATGCAAGGAACAGTGAATTGGAGTTCG
TCTTGTTATAATTAGCTTCTTGGGGTATCTTTAAATACTGTAGAAAAGAGGAAGGAAATAATAAATGGCTAAAATGAGAATATCACCGGA
ATTGAAAAAACTGATCGAAAAATACCGCTGCGTAAAAGATACGGAAGGAATGTCTCCTGCTAAGGTATATAAGCTGGTGGGAGAAAATGA
AAACCTATATTTAAAAATGACGGACAGCCGGTATAAAGGGACCACCTATGATGTGGAACGGGAAAAGGACATGATGCTATGGCTGGAAGG
AAAGCTGCCTGTTCCAAAGGTCCTGCACTTTGAACGGCATGATGGCTGGAGCAATCTGCTCATGAGTGAGGCCGATGGCGTCCTTTGCTC
GGAAGAGTATGAAGATGAACAAAGCCCTGAAAAGATTATCGAGCTGTATGCGGAGTGCATCAGGCTCTTTCACTCCATCGACATATCGGA
TTGTCCCTATACGAATAGCTTAGACAGCCGCTTAGCCGAATTGGATTACTTACTGAATAACGATCTGGCCGATGTGGATTGCGAAAACTG
GGAAGAAGACACTCCATTTAAAGATCCGCGCGAGCTGTATGATTTTTTAAAGACGGAAAAGCCCGAAGAGGAACTTGTCTTTTCCCACGG
CGACCTGGGAGACAGCAACATCTTTGTGAAAGATGGCAAAGTAAGTGGCTTTATTGATCTTGGGAGAAGCGGCAGGGCGGACAAGTGGTA
TGACATTGCCTTCTGCGTCCGGTCGATCAGGGAGGATATCGGGGAAGAACAGTATGTCGAGCTATTTTTTGACTTACTGGGGATCAAGCC
TGATTGGGAGAAAATAAAATATTATATTTTACTGGATGAATTGTTTTAGTACCTAGAATGCATGACCAAAATCCCTTAACGTGAGTTTTC
GTTCCACTGAGCGTCAGACCCCGTAGAAAAGATCAAAGGATCTTCTTGAGATCCTTTTTTTCTGCGCGTAATCTGCTGCTTGCAAACAAA
AAAACCACCGCTACCAGCGGTGGTTTGTTTGCCGGATCAAGAGCTACCAACTCTTTTTCCGAAGGTAACTGGCTTCAGCAGAGCGCAGAT
ACCAAATACTGTCCTTCTAGTGTAGCCGTAGTTAGGCCACCACTTCAAGAACTCTGTAGCACCGCCTACATACCTCGCTCTGCTAATCCT
GTTACCAGTGGCTGCTGCCAGTGGCGATAAGTCGTGTCTTACCGGGTTGGACTCAAGACGATAGTTACCGGATAAGGCGCAGCGGTCGGG
CTGAACGGGGGGTTCGTGCACACAGCCCAGCTTGGAGCGAACGACCTACACCGAACTGAGATACCTACAGCGTGAGCTATGAGAAAGCGC
CACGCTTCCCGAAGGGAGAAAGGCGGACAGGTATCCGGTAAGCGGCAGGGTCGGAACAGGAGAGCGCACGAGGGAGCTTCCA GGGGGAAA
CGCCTGGTATCTTTATAGTCCTGTCGGGTTTCGCCACCTCTGACTTGAGCGTCGATTTTTGTGATGCTCGTCAGGGGGGCGGAGCCTATG
GAAAAACGCCAGCAACGCGGCCTTTTTACGGTTCCTGGCCTTTTGCTGGCCTTTTGCTCACATGTTCTTTCCTGCGTTATCCCCTGATTC
TGTGGATAACCGTATTACCGCCTTTGAGTGAGCTGATACCGCTCGCCGCAGCCGAACGACCGAGCGCAGCGAGTCAGTGAGCGAGGAAGC
GGAAGAGCGCCTGATGCGGTATTTTCTCCTTACGCATCTGTGCGGTATTTCACACCGCATATGGTGCACTCTCAGTACAATCTGCTCTGA
TGCCGCATAGTTAAGCCAGTATACACTCCGCTATCGCTACGTGACTGGGTCATGGCTGCGCCCCGACACCCGCCAACACCCGCTGACGCG
CCCTGACGGGCTTGTCTGCTCCCGGCATCCGCTTACAGACAAGCTGTGACCGTCTCCGGGAGCTGCATGTGTCAGAGGTTTTCACCGTCA
TCACCGAAACGCGCGAGGCAGGGTGCCTTGATGTGGGCGCCGGCGGTCGAGTGGCGACGGCGCGGCTTGTCCGCGCCCTGGTAGATTGCC
TGGCCGTAGGCCAGCCATTTTTGAGCGGCCAGCGGCCGCGATAGGCCGACGCGAAGCGGCGGGGCGTAGGGAGCGCAGCGACCGAAGGGT
AGGCGCTTTTTGCAGCTCTTCGGCTGTGCGCTGGCCAGACAGTTATGCACAGGCCAGGCGGGTTTTAAGAGTTTTAATAAGTTTTAAAGA
GTTTTAGGCGGAAAAATCGCCTTTTTTCTCTTTTATATCAGTCACTTACATGTGTGACCGGTTCCCAATGTACGGCTTTGGGTTCCCAAT
GTACGGGTTCCGGTTCCCAATGTACGGCTTTGGGTTCCCAATGTACGTGCTATCCACAGGAAAGAGAACTTTTCGACCTTTTTCCCCTGC
TAGGGCAATTTGCCCTAGCATCTGCTCCGTACATTAGGAACCGGCGGATGCTTCGCCCTCGATCAGGTTGCGGTAGCGCATGACTAGGAT
CGGGCCAGCCTGCCCCGCCTCCTCCTTCAAATCGTACTCCGGCAGGTCATTTGACCCGATCAGCTTGCGCACGGTGAAACAGAACTTCTT
GAACTCTCCGGCGCTGCCACTGCGTTCGTAGATCGTCTTGAACAACCATCTGGCTTCTGCCTTGCCTGCGGCGCGGCGTGCCAGGCGGTA
GAGAAAACGGCCGATGCCGGGATCGATCAAAAAGTAATCGGGGTGAACCGTCAGCACGTCCGGGTTCTTGCCTTCTGTGATCTCGCGGTA
CATCCAATCAGCTAGCTCGATCTCGATGTACTCCGGCCGCCCGGTTTCGCTCTTTACGATCTTGTAGCGGCTAATCAAGGCTTCACCCTC
GGATACCGTCACCAGGCGGCCGTTCTTGGCCTTCTTCGTACGCTGCATGGCAACGTGCGTGGTGTTTAACCGAATGCAGGTTTCTACCAG
GTCGTCTTTCTGCTTTCCGCCATCGGCTCGCCGGCAGAACTTGAGTACGTCCGCAACGTGTGGACGGAACACGCGGCCGGGCTTGTCTCC
CTTCCCTTCCCGGTATCGGTTCATGGATTCGGTTAGATGGGAAACCGCCATCAGTACCAGGTCGTAATCCCACACACTGGCCATGCCGGC
CGGCCCTGCGGAAACCTCTACGTGCCCGTCTGGAAGCTCGTAGCGGATCACCTCGCCAGCTCGTCGGTCACGCTTCGACAGACGGAAAAC
GGCCACGTCCATGATGCTGCGACTATCGCGGGTGCCCACGTCATAGAGCATCGGAACGAAAAAATCTGGTTGCTCGTCGCCCTTGGGCGG
CTTCCTAATCGACGGCGCACCGGCTGCCGGCGGTTGCCGGGATTCTTTGCGGATTCGATCAGCGGCCGCTTGCCACGATTCACCGGGGCG
TGCTTCTGCCTCGATGCGTTGCCGCTGGGCGGCCTGCGCGGCCTTCAACTTCTCCACCAGGTCATCACCCAGCGCCGCGCCGATTTGTAC
CGGGCCGGATGGTTTGCGACCGTCACGCCGATTCCTCGGGCTTGGGGGTTCCAGTGCCATTGCAGGGCCGGCAGACAACCCAGCCGCTTA
CGCCTGGCCAACCGCCCGTTCCTCCACACATGGGGCATTCCACGGCGTCGGTGCCTGGTTGTTCTTGATTTTCCATGCCGCCTCCTTTAG
CCGCTAAAATTCATCTACTCATTTATTCATTTGCTCATTTACTCTGGTAGCTGCGCGATGTATTCAGATAGCAGCTCGGTAATGGTCTTG
CCTTGGCGTACCGCGTACATCTTCAGCTTGGTGTGATCCTCCGCCGGCAACTGAAAGTTGACCCGCTTCATGGCTGGCGTGTCTGCCAGG
CTGGCCAACGTTGCAGCCTTGCTGCTGCGTGCGCTCGGACGGCCGGCACTTAGCGTGTTTGTGCTTTTGCTCATTTTCTCTTTACCTCAT
TAACTCAAATGAGTTTTGATTTAATTTCAGCGGCCAGCGCCTGGACCTCGCGGGCAGCGTCGCCCTCGGGTTCTGATTCAAGAACGGTTG
TGCCGGCGGCGGCAGTGCCTGGGTAGCTCACGCGCTGCGTGATACGGGACTCAAGAATGGGCAGCTCGTACCCGGCCAGCGCCTCGGCAA
CCTCACCGCCGATGCGCGTGCCTTTGATCGCCCGCGACACGACAAAGGCCGCTTGTAGCCTTCCATCCGTGACCTCAATGCGCTGCTTAA
CCAGCTCCACCAGGTCGGCGGTGGCCCATATGTCGTAAGGGCTTGGCTGCACCGGAATCAGCACGAAGTCGGCTGCCTTGATCGCGGACA
CAGCCAAGTCCGCCGCCTGGGGCGCTCCGTCGATCACTACGAAGTCGCGCCGGCCGATGGCCTTCACGTCGCGGTCAATCGTCGGGCGGT
CGATGCCGACAACGGTTAGCGGTTGATCTTCCCGCACGGCCGCCCAATCGCGGGCACTGCCCTGGGGATCGGAATCGACTAACAGAACAT
CGGCCCCGGCGAGTTGCAGGGCGCGGGCTAGATGGGTTGCGATGGTCGTCTTGCCTGACCCGCCTTTCTGGTTAAGTACAGCGATAACCT
TCATGCGTTCCCCTTGCGTATTTGTTTATTTACTCATCGCATCATATACGCAGCGACCGCATGACGCAAGCTGTTTTACTCAAATACACA
TCACCTTTTTAGACGGCGGCGCTCGGTTTCTTCAGCGGCCAAGCTGGCCGGCCAGGCCGCCAGCTTGGCATCAGACAAAC CGGCCAGGAT
TTCATGCAGCCGCACGGTTGAGACGTGCGCGGGCGGCTCGAACACGTACCCGGCCGCGATCATCTCCGCCTCGATCTCTTCGGTAATGAA
AAACGGTTCGTCCTGGCCGTCCTGGTGCGGTTTCATGCTTGTTCCTCTTGGCGTTCATTCTCGGCGGCCGCCAGGGCGTCGGCCTCGGTC
AATGCGTCCTCACGGAAGGCACCGCGCCGCCTGGCCTCGGTGGGCGTCACTTCCTCGCTGCGCTCAAGTGCGCGGTACAGGGTCGAGCGA
TGCACGCCAAGCAGTGCAGCCGCCTCTTTCACGGTGCGGCCTTCCTGGTCGATCAGCTCGCGGGCGTGCGCGATCTGTGCCGGGGTGAGG
GTAGGGCGGGGGCCAAACTTCACGCCTCGGGCCTTGGCGGCCTCGCGCCCGCTCCGGGTGCGGTCGATGATTAGGGAACGCTCGAACTCG

GCAATGCCGGCGAACACGGTCAACACCATGCGGCCGGCCGGCGTGGTGGTGTCGGCCCACGGCTCTGCCAGGCTACGCAGGCCCGCGCCG
GCCTCCTGGATGCGCTCGGCAATGTCCAGTAGGTCGCGGGTGCTGCGGGCCAGGCGGTCTAGCCTGGTCACTGTCACAACGTCGCCAGGG
CGTAGGTGGTCAAGCATCCTGGCCAGCTCCGGGCGGTCGCGCCTGGTGCCGGTGATCTTCTCGGAAAACAGCTTGGTGCAGCCGGCCGCG
TGCAGTTCGGCCCGTTGGTTGGTCAAGTCCTGGTCGTCGGTGCTGACGCGGGCATAGCCCAGCAGGCCAGCGGCGGCGCTCTTGTTCATG
GCGTAATGTCTCCGGTTCTAGTCGCAAGTATTCTACTTTATGCGACTAAAACACGCGACAAGAAAACGCCAGGAAAAGGGCAGGGCGGCA
GCCTGTCGCGTAACTTAGGACTTGTGCGACATGTCGTTTTCAGAAGACGGCTGCACTGAACGTCAGAAGCCGACTGCACTATAGCAGCGG
AGGGGTTGGATCAAAGTACTTTGATCCCGAGGGGAACCCTGTGGTTGGCATGCACATACAAATGGACGAACGGATAAACCTTTTCACGCC
CTTTTAAATATCCGTTATTCTAATAAACGCTCTTTTCTCTTAGGTTTACCCGCCAATATATCCTGTCAAACACTGATAGTTTAAACTGAA
GGCGGGAAACGACAATCTGATCCAAGCTCAAGCTGCTCTAGCATTCGCCATTCAGGCTGCGCAACTGTTGGGAAGGGCGATCGGTGCGGG
CCTCTTCGCTATTACGCCAGCTGGCGAAAGGGGGATGTGCTGCAAGGCGATTAAGTTGGGTAACGCCAGGGTTTTCCCAGTCACGACGTT
GTAAAACGACGGCCAGTGCCAAGCTTGGCGTGCCTGCAGGTCAACATGGTGGAGCACGACACACTTGTCTACTCCAAAAATATCAAAGAT
ACAGTCTCAGAAGACCAAAGGGCAATTGAGACTTTTCAACAAAGGGTAATATCCGGAAACCTCCTCGGATTCCATTGCCCAGCTATCTGT
CACTTTATTGTGAAGATAGTGGAAAAGGAAGGTGGCTCCTACAAATGCCATCATTGCGATAAAGGAAAGGCCATCGTTGAAGATGCCTCT
GCCGACAGTGGTCCCAAAGATGGACCCCCACCCACGAGGAGCATCGTGGAAAAAGAAGACGTTCCAACCACGTCTTCAAAGCAAGTGGAT
TGATGTGATAACATGGTGGAGCACGACACACTTGTCTACTCCAAAAATATCAAAGATACAGTCTCAGAAGACCAAAGGGCAATTGAGACT
TTTCAACAAAGGGTAATATCCGGAAACCTCCTCGGATTCCATTGCCCAGCTATCTGTCACTTTATTGTGAAGATAGTGGAAAAGGAAGGT
GGCTCCTACAAATGCCATCATTGCGATAAAGGAAAGGCCATCGTTGAAGATGCCTCTGCCGACAGTGGTCCCAAAGATGGACCCCCACCC
ACGAGGAGCATCGTGGAAAAAGAAGACGTTCCAACCACGTCTTCAAAGCAAGTGGATTGATGTGATATCTCCACTGACGTAAGGGATGAC
GCACAATCCCACTATCCTTCGCAAGACCCTTCCTCTATATAAGGAAGTTCATTTCATTTGGAGAGGACCTCGACTCTAGAGGATCCCCGG
GTACCGGGCCCCCCCTCGAGGCGCGCCAAGCTATCAAACAAGTTTGTACAAAAAAGCAGGCTCCGCGGCCGCCCCCTTCACCTGTA**GGCA
TGGGCGGTGTAGGCAAGATAGACCATTTA**AGAGACCATTAGGCACCCCAGGCTTTACACTTTATGCTTCCGGCTCGTATAATGTGTGGAT
TTTGAGTTAGGAGCCGTCGAGATTTTCAGGAGCTAAGGAAGCTAAAATGGAGAAAAAAATCACTGGATATACCACCGTTGATATATCCCA
ATGGCATCGTAAAGAACATTTTGAGGCATTTCAGTCAGTTGCTCAATGTACCTATAACCAGACCGTTCAGCTGGATATTACGGCCTTTTT
AAAGACCGTAAAGAAAAATAAGCACAAGTTTTATCCGGCCTTTATTCACATTCTTGCCCGCCTGATGAATGCTCATCCGGAGTTCCGTAT
GGCAATGAAAGACGGTGAGCTGGTGATATGGGATAGTGTTCACCCTTGTTACACCGTTTTCCATGAGCAAACTGAAACGTTTTCATCGCT
CTGGAGTGAATACCACGACGATTTCCGGCAGTTTCTACACATATATTCGCAAGATGTGGCGTGTTACGGTGAAAACCTGGCCTATTTCCC
TAAAGGGTTTATTGAGAATATGTTTTTCGTCTCAGCCAATCCCTGGGTGAGTTTCACCAGTTTTGATTTAAACGTGGCCAATATGGACAA
CTTCTTCGCCCCCGTTTTCACCATGGGCAAATATTATACGCAAGGCGACAAGGTGCTGATGCCGCTGGCGATTCAGGTTCATCATGCCGT
TTGTGATGGCTTCCATGTCGGCAGAATGCTTAATGAATTACAACAGTACTGCGATGAGTGGCAGGGCGGGGCGTAAACGCGTGGAGCCGG
CTTACTAAAAGCCAGATAACAGTATGCGTATTTGCGCGCTGATTTTTGCGGTATAAGAATATATACTGATATGTATACCCGAAGTATGTC
AAAAAGAGGTATGCTATGAAGCAGCGTATTACAGTGACAGTTGACAGCGACAGCTATCAGTTGCTCAAGGCATATATGATGTCAATATCT
CCGGTCTGGTAAGCACAACCATGCAGAATGAAGCCCGTCGTCTGCGTGCCGAACGCTGGAAAGCGGAAAATCAGGAAGGGATGGCTGAGG
TCGCCCGGTTTATTGAAATGAACGGCTCTTTTGCTGACGAGAACAGGGGCTGGTGAAATGCAGTTTAAGGTTTACACCTATAAAAGAGAG
AGCCGTTATCGTCTGTTTGTGGATGTACAGAGTGATATTATTGACACGCCCGGGCCGACGGATGGTGATCCCCCTGGCCAGTGCACGTCTG
CTGTCAGATAAAGTCTCCCGTGAACTTTACCCGGTGGTGCATATCGGGGATGAAAGCTGGCGCATGATGACCACCGATATGGCCAGTGTG
CCGGTTTCCGTTATCGGGGAAGAAGTGGCTGATCTCAGCCACCGCGAAAATGACATCAAAAACGCCATTAACCTGATGTTCTGGGGAATA
TAAATGTCAGGCTCCCTTATACACAGCCAGTCTGCACCTCGACGGTCTCACATTAAGGGTGGGCGCGCCGACCCAGCTTTCTTGTACAAA
GTGGTTCGATAATTCCTTAATTAACTAGTTCTAGAGCGGCCGCCCACCGCGGTGGAGCTCGAATTTCCCCGATCGTTCAAACATTTGGCA
ATAAAGTTTCTTAAGATTGAATCCTGTTGCCGGTCTTGCGATGATTATCATATAATTTCTGTTGAATTACGTTAAGCATGTAATAATTAA
CATGTAATGCATGACGTTATTTATGAGATGGGTTTTTATGATTAGAGTCCCGCAATTATACATTTAATACGCGATAGAAAACAAAATATA
GCGCGCAAACTAGGATAAATTATCGCGCGCGGTGTCATCTATGTTACTGAATTCGTAATCATGGTCATAGCTGTTTCCTGTGTGAAATTG
TTATCCGCTCACAATTCCACACAACATACGAGCCGGAAGCATAAAGTGTAAAGCCTGGGGTGCCTAATGAGTGAGCTAACTCACATTAAT
TGCGTTGCGCTCACTGCCCGCTTTCCAGTCGGGAAACCTGTCGTGCCAGCTGCATTAATGAATCGGCCAACGCGCGGGGAGAGGCGGTTT
GCGTATTGGCTAGAGCAGCTTGCCAACATGGTGGAGCACGACACTCTCGTCTACTCCAAGAATATCAAAGATACAGTCTCAGAAGACCAA
AGGGCTATTGAGACTTTTCAACAAAGGGTAATATCGGGAAACCTCCTCGGATTCCATTGCCCAGCTATCTGTCACTTCATCAAAAGGACA
GTAGAAAAGGAAGGTGGCACCTACAAATGCCATCATTGCGATAAAGGAAAGGCTATCGTTCAAGATGCCTCTGCCGACAGTGGTCCCAAA
GATGGACCCCCACCCACGAGGAGCATCGTGGAAAAAGAAGACGTTCCAACCACGTCTTCAAAGCAAGTGGATTGATGTGATAACATGGTG
GAGCACGACACTCTCGTCTACTCCAAGAATATCAAAGATACAGTCTCAGAAGACCAAAGGGCTATTGAGACTTTTCAACAAAGGGTAATA
TCGGGAAACCTCCTCGGATTCCATTGCCCAGCTATCTGTCACTTCATCAAAAGGACAGTAGAAAAGGAAGGTGGCACCTACAAATGCCAT
CATTGCGATAAAGGAAAGGCTATCGTTCAAGATGCCTCTGCCGACAGTGGTCCCAAAGATGGACCCCCACCCACGAGGAGCATCGTGGAA
AAAGAAGACGTTCCAACCACGTCTTCAAAGCAAGTGGATTGATGTGATATCTCCACTGACGTAAGGGATGACGCACAATCCCACTATCCT
TCGCAAGACCTTCCTCTATATAAGGAAGTTCATTTCATTTGGAGAGGACACGCTGAAATCACCAGTCTCTCTCTACAAATCTATCTCTCT
CGAGCTTTCGCAGATCCCGGGGGGCAATGAGATATGAAAAAGCCTGAACTCACCGCGACGTCTGTCGAGAAGTTTCTGATCGAAAAGTTC
GACAGCGTCTCCGACCTGATGCAGCTCTCGGAGGGCGAAGAATCTCGTGCTTTCAGCTTCGATGTAGGAGGGCGTGGATATGTCCTGCGG
GTAAATAGCTGCGCCGATGGTTTCTACAAAGATCGTTATGTTTATCGGCACTTTGCATCGGCCGCGCTCCCGATTCCGGAAGTGCTTGAC
ATTGGGGAGTTTAGCGAGAGCCTGACCTATTGCATCTCCCGCCGTGCACAGGGTGTCACGTTGCAAGACCTGCCTGAAACCGAACTGCCC
GCTGTTCTACAACCGGTCGCGGAGGCTATGGATGCGATCGCTGCGGCCGATCTTAGCCAGACGAGCGGGTTCGGCCCATTCGGACCGCAA
GGAATCGGTCAATACACTACATGGCGTGATTTCATATGCGCGATTGCTGATCCCCATGTGTATCACTGGCAAACTGTGATGGACGACACC
GTCAGTGCGTCCGTCGCGCAGGCTCTCGATGAGCTGATGCTTTGGGCCGAGGACTGCCCCGAAGTCCGGCACCTCGTGCACGCGGATTTC
GGCTCCAACAATGTCCTGACGGACAATGGCCGCATAACAGCGGTCATTGACTGGAGCGAGGCGATGTTCGGGGATTCCCAATACGAGGTC
GCCAACATCTTCTTCTGGAGGCCGTGGTTGGCTTGTATGGAGCAGCAGACGCGCTACTTCGAGCGGAGGCATCCGGAGCTTGCAGGATCG
CCACGACTCCGGGCGTATATGCTCCGCATTGGTCTTGACCAACTCTATCAGAGCTTGGTTGACGGCAATTTCGATGATGCAGCTTGGGCG
CAGGGTCGATGCGACGCAATCGTCCGATCCGGAGCCGGGACTGTCGGGCGTACACAAATCGCCCGCAGAAGCGCGGCCGTCTGGACCGAT

GGCTGTGTAGAAGTACTCGCCGATAGTGGAAACCGACGCCCCAGCACTCGTCCGAGGGCAAAGAAATAGAGTAGATGCCGACCGGATCTG
TCGATCGACAAGCTCGAGTTTCTCCATAATAATGTGTGAGTAGTTCCCAGATAAGGGAATTAGGGTTCCTATAGGGTTTCGCTCATGTGT
TGAGCATATAAGAAACCCTTAGTATGTATTTGTATTTGTAAAATACTTCTATCAATAAAATTTCTAATTCCTAAAACCAAAATCCAGTAC
TAAAATCCAGATCCCCCGAATTAATTCGGCGTTAATTCAGTACATTAAAAACGTCCGCAATGTGTTATTAAGTTGTCTAAGCGTCAATTT
GTTTACACCACAATATATCCTGCCA

**SlmiR482b target site**

***AtTAS1c*-derived spacer**

T-DNA right border

T-DNA left border

*ccd*B gene

*Bsa*I site

Inverted *Bsa*I site

Chloramphenicol resistance gene

attB1

attB2

Nos terminator

CaMV promoter

kanamycin resistance gene

Hygromycin resistance gene

2x35S CaMV promoter

CaMV terminator
